# Mosaic chromosomal alterations in blood across ancestries via whole-genome sequencing

**DOI:** 10.1101/2022.11.07.515222

**Authors:** Yasminka A. Jakubek, Ying Zhou, Adrienne Stilp, Jason Bacon, Justin Wong, Zuhal Ozcan, Donna Arnett, Kathleen Barnes, Josh Bis, Eric Boerwinkle, April Carson, Daniel Chasman, Michael Cho, Matthew P. Conomos, Nancy Cox, Margaret Doyle, Myriam Fornage, Xiuqing Guo, Sharon Kardia, Joshua P. Lewis, Ruth J. Loos, Xiaolong Ma, Mitchell Machiela, Taralynn M. Mack, Rasika Mathias, Braxton D. Mitchell, Kari North, Nathan Pankratz, Patricia Peyser, Michael H. Preuss, Bruce Psaty, Laura M. Raffield, Ramachandran S. Vasan, Susan Redline, Stephen S. Rich, Jerome I. Rotter, Edwin Silverman, Jennifer Smith, Margaret Taub, Jeong Yun, Yun Li, Pinkal Desai, Alexander G. Bick, Alexander P. Reiner, Paul Scheet, Paul L. Auer, the NHLBI TOPMed program

## Abstract

Mosaic mutations in blood are common with increasing age and are prognostic markers for cancer, cardiovascular dysfunction and other diseases. This group of acquired mutations include megabase-scale mosaic chromosomal alterations (mCAs). These large mutations have mainly been surveyed using SNP array data from individuals of European (EA) or Japanese genetic ancestry. To gain a better understanding of mCA rates and associated risk factors in genetically diverse populations, we surveyed whole genome sequencing data from 67,390 individuals, including 20,132 individuals of African ancestry (AA), and 7,608 of Hispanic ancestry (HA) with deep (30X) whole genome sequencing data from the NHLBI Trans Omics for Precision Medicine (TOPMed) program. We adapted an existing mCA calling algorithm for application to WGS data, and observed higher sensitivity with WGS data, compared with array-based data, in uncovering mCAs at low mutant cell fractions. As in previous reports, we observed a strong association with age and a non-uniform distribution of mCAs across the genome. The presence of autosomal (but not chromosome X) mCAs was associated with an increased risk of both lymphoid and myeloid malignancies. After adjusting for age, we found that individuals of European ancestry have the highest rates of autosomal mCAs, mirroring the higher rate of leukemia in this group. Our analysis also uncovered higher rates of chromosome X mCAs in AA and HA compared to EA, again after adjusting for age. Germline variants in *ATM* and *MPL* showed strong associations with mCAs in *cis*, including ancestry specific variants. And rare variant gene-burden analysis confirmed the association of putatively protein altering variants in *ATM* and *MPL* with mCAs in *cis*. Individual rare variants in *DCPS, ADM17, PPP1R16B*, and *TET2* were all associated with autosomal mCAs and rare variants in *OR4C16* were associated with chromosome X mCAs in females. There was significant enrichment of co-occurrence of CHIP mutations and mCAs both altering cancer associated genes *TET2, DNMT3A, JAK2, CUX1*, and *TP53*. Overall, our study demonstrates that rates of mCAs differ across populations and that rare inherited germline variants are strongly associated with mCAs across genetically diverse populations. These results strongly motivate further studies of mCAs in under-represented populations to better understand the causes and consequences of this class of somatic variation.

## INTRODUCTION

Mosaicism refers to the presence of genetically distinct lineages of cells resulting from a single zygote in a multicellular organism. The clone with a somatic mutation may comprise a substantial fraction of cells in a tissue, which have risen to detectable frequency due to selective advantage or drift. Surveys of blood samples from healthy donors have revealed extensive age-related clonal mosaicism, which can involve somatic mutations ranging in size from a single nucleotide to large, typically megabase-scale alterations spanning an entire chromosome.^1-7^ Such studies have additionally demonstrated that the presence of these acquired mutations in autosomes are more common in men and confer a ∼10-fold higher risk for the development of hematological malignancies in otherwise healthy adults.^3,4,8,9^ Specifically these studies have identified large (>1-2 Mb) mosaic chromosomal alterations (mCAs) which include chromosomal losses, gains, and copy neutral loss of heterozygosity (CN-LOH). Other studies have surveyed mosaic somatic single nucleotide variants (SNVs) in the blood. In particular, mosaicism for acquired leukemogenic SNV mutations in individuals without evidence of hematologic malignancy, dysplasia, or cytopenia is known as clonal hematopoiesis of indeterminate potential (CHIP). These studies of mosaic SNVs have shown an ∼13-fold higher risk for development of hematological malignancies in those with CHIP mutations.^10-12^ Beyond cancer, mosaicism in blood has been associated with other chronic diseases, further highlighting its potential as an endophenotype to understand etiology or as a biomarker with clinical utility.^11,13-15^

To date, the largest studies of mCAs have surveyed DNA array data from individuals of European or Japanese genetic ancestries.^6,16^ Yet, the landscape of mCAs in more genetically diverse populations and from whole genome sequencing (WGS) remains underexplored. To address this void we investigated mCAs using WGS data from the Trans-Omics for Precision Medicine (TOPMed) program, in 67,390 individuals, including 20,132 individuals of African American ancestry (AA), 7,608 individuals of Hispanic ancestry (HA), and 1,203 individuals of East Asian (EAS) ancestry. We demonstrate for the first time the use of a haplotype-based methodology for the detection of mCAs from high coverage (30X) WGS data. This methodology allowed for sensitive detection of mCAs at mutant cell fractions below 1% and enabled association analyses of both rare and common germline variation with the presence of mCAs.

## RESULTS

### Genomic landscape of mCAs

We detected 3,659 autosomal mCAs in 67,390 TOPMed samples (**Figure 1 & Table S1**). A total of 3,017 samples (4.47%) had at least one detectable autosomal mCA, of which 0.49% had mCAs in more than one autosomal chromosome. As reported previously, the rate of mCAs increased with age (**Figure 2A**), and the rate of autosomal mCAs for males was higher than that for females (age-adjusted OR = 1.19, P = 6.3e-05, **Figure 2A**). To investigate the accuracy of our mCA calls, we compared the mCA detection in our WGS data to array-based mCA calls on a subset of individuals. Overall, we found that mCA detection was more sensitive with the TOPMed WGS data, particularly for mCAs with clonal fractions below 3% (see **Supplement**).

**Figure 1.**
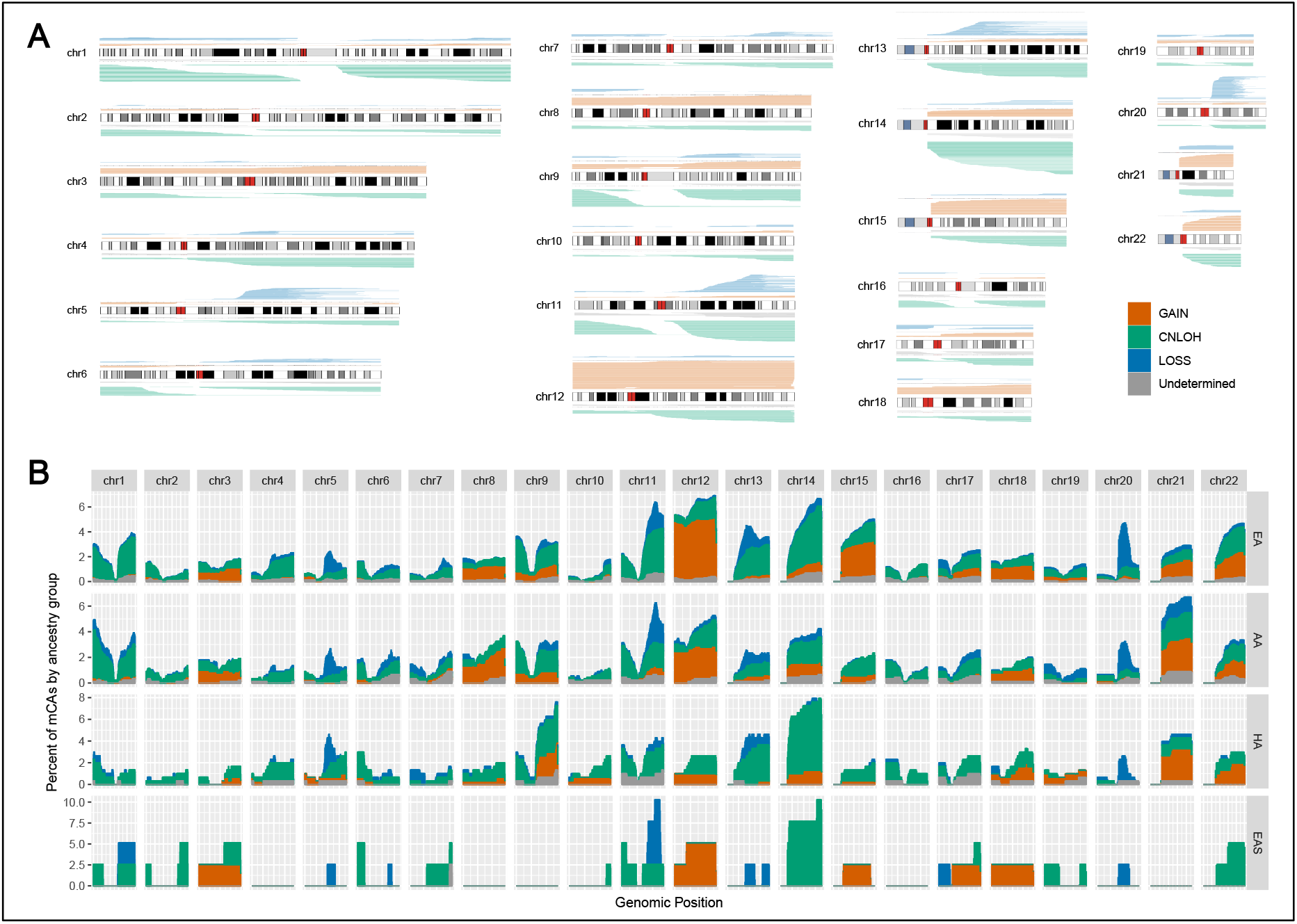
Genomic distribution of autosomal mCAs. **A)** mCA calls across autosomal chromosomes **B)** Histogram of mCA calls across the genome for each genetic ancestry group. **B)** X-axis are 1Mb windows for each chromosome and Y-axis is the percent of mCA calls for a given genetic ancestry group that span the genomic window.

**Figure 2.**
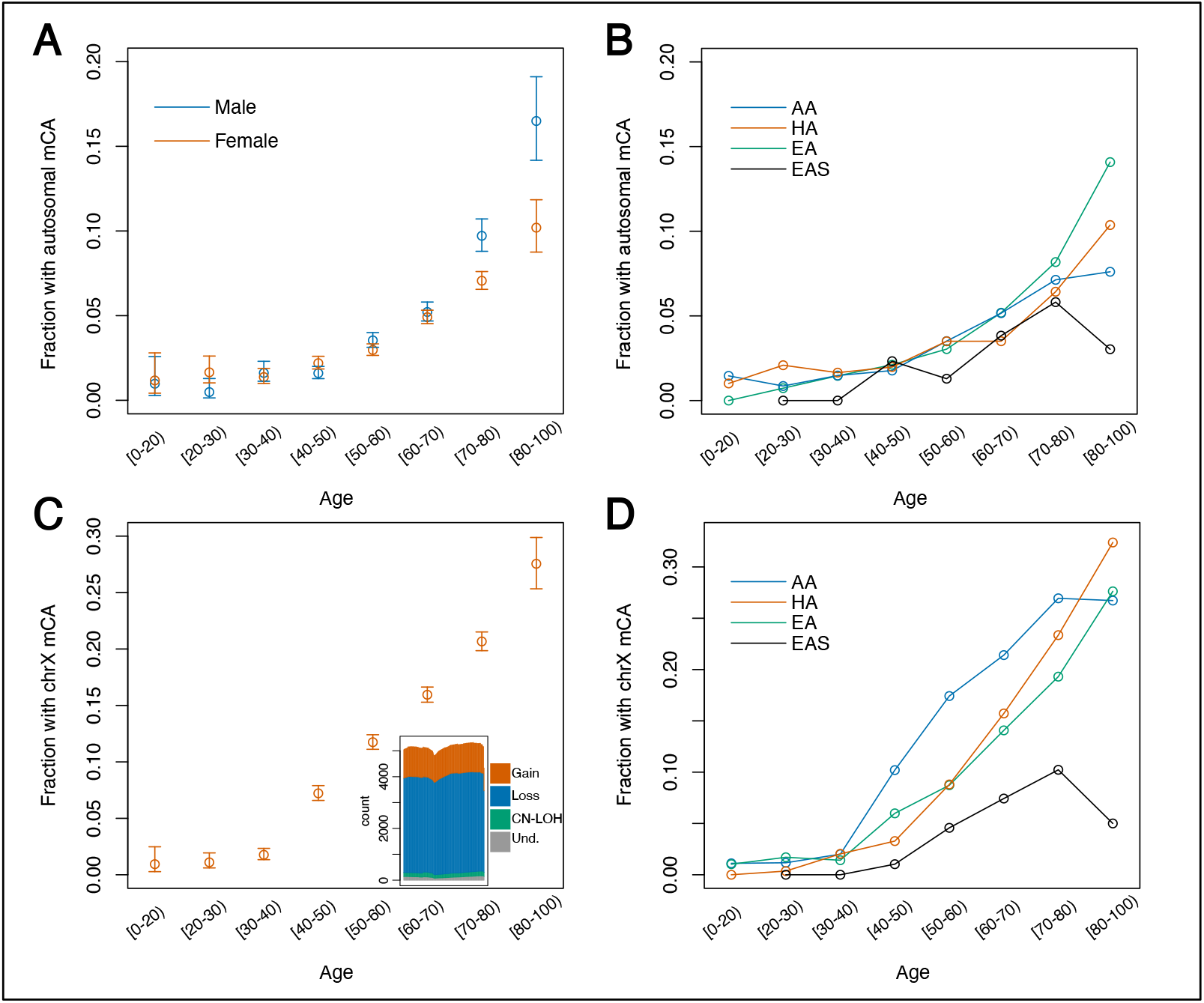
Rate mCAs by age. **A)** Fraction of females and males with one or more autosomal mCA across age bins. **B)** Fraction of individuals across different genetic ancestry groups with one or more autosomal mCA across age bins. **C)** Fraction of females with a chromosome X mCA across age bins. Histogram of chromosome X mCA calls. X-axis are 1Mb windows across chromosome X and Y-axis is the number of mCA calls that span the genomic window. **D)** Fraction of females across different genetic ancestry groups with a chromosome X mCA across age bins.

Autosomal mCAs were categorized as gain, loss, CN-LOH, or undetermined. The most frequent autosomal mCAs (n > 100) were 14q CN-LOH, 12p and 12q gains, 20q loss, 11q CN-LOH, and 1p CN-LOH (**Figure 1 & Table S2**). We tested for differences in the rates of each autosomal mCA across chromosome arms in males and females. We found significant enrichment in males for chromosome 20q arm loss (OR = 2.76, P = 1.1e-05) and 15q gain (OR = 2.73, P = 2.1e-03) after adjustment for multiple testing. Multiple other loci exhibiting mCAs had a significant sex-specific enrichment prior to multiple testing correction (n = 11) with all having higher rates in males (**Table S3**).

The majority (82%) of detectable autosomal mCAs were estimated to be present at cell fractions less than 10%, with a large proportion (44%) present at estimated cell fractions less than 3% (**Sup. Figure 1A**). Only 8.5% of autosomal mCAs were present at estimated cell fractions greater than 20%. When restricting to mCAs present at cell fractions of 10% or greater, chromosome 20 mCAs were most frequent (∼2% of all autosomal mCAs with CF > 10%), followed by mCAs on chromosome 12 (∼1% of all autosomal mCAs with CF > 10%). This distribution contrasts with the distribution of autosomal mCAs across all cell fractions, where the most frequently altered chromosomes are 11, 12, and 14 (**Sup. Figure 2**). This variation suggests differential fitness advantages across clones with different mCAs.

Using the same methodology as for autosomal mCAs, we surveyed mCAs on chromosome X (chrX) from 41,895 female samples in the TOPMed cohort and identified 6,207 mCAs. The rate of chrX mCAs was significantly higher (P=2.2e-16) than for autosomal mCAs with 13.7% of females harboring an mCA on chrX in contrast to 4.38% with mCAs on the autosomes **(Sup. Figure 4)**. Overall mCAs on chrX had lower estimated cell fraction than mCAs on the autosomes (**Sup. Fig 1**). Most of the mCAs on chrX (68.8%) were losses (**Figure 2C & Sup. Figure 5**). The presence of an mCA on chrX showed a positive association with autosomal mCAs (OR=1.21, p = 0.003; adjusted for age).

### Distribution of mCAs across ancestries

TOPMed samples have been previously categorized into five genetic ancestry groups, namely AA, HA, EA, EAS, and a fifth set of individuals for which genetic ancestry could not be confidently determined (see **Methods** and **Table S1**). We sought to make comparisons of mCA rates across these ancestry groups. Because haplotype-based detection of mCAs is made possible by analysis of intermediate-level (within-sample) allele-specific data (i.e., “B allele” frequencies) *at heterozygous genotypes*, the higher heterozygosity levels in HA, EA, and AA relative to EAS could drive differences in sensitivity for detection of mCAs (**Table S4**). To control for the impact of variable heterozygosity rates, we down-sampled heterozygous sites in the AA, HA, and EA groups to match the distribution in the EAS group which has the lowest heterozygosity rate and re-ran the mCA detection procedure (**see Methods**). After down-sampling, the AA, HA, and EAS groups exhibited significantly lower autosomal mCA rates relative to EA (OR = 0.76, P = 1.2e-05 for AA; OR = 0.71, P = 0.0009 for HA; OR = 0.62, P = 0.007 for EAS, adjusted for age, age^2^, sex and study cohort).

As with autosomal mCAs, we contrasted chrX mCA rates across genetic ancestries by down-sampling heterozygous sites on chrX in the AA, EAS, and HA groups to match the EAS distribution as well. After this adjustment we observed a higher chrX mCA rate in AA and HA compared to EA (OR = 1.67, P = 2.5e-33 for AA & OR = 1.36, P = 0.00013 for HA). The chrX mCA rate for EAS was lower compared to EA (OR = 0.49, P = 3.2e-5). To determine if the association was driven by a specific type of mCA we tested for associations between genetic ancestry and chrX loss and gain separately. ChrX loss rates were higher in AA (OR = 1.59, P = 2.58e-20) and in HA (OR = 1.42, P = 2.13e-05) compared to EA ancestry groups. As with autosomal mCAs, chrX loss was lower in EAS (OR = 0.41, P = 3.5e-07) compared to EA and the AA group demonstrated a higher rate of chrX gains compared to the EA group (OR = 2.06, P = 6.27e-22).

Next, we contrasted the autosomal regions that harbored mCAs between AA and EA, the two genetic ancestry groups with the largest sample size and thus most amenable to such a comparison (**Figure 1B & Sup. Figure 3**). In the EA group, autosomal mCAs were observed most frequently on chromosomes 11, 12, and 14. For the AA group, the most frequent autosomal mCAs were observed on chromosomes 11, 12, and 21. When mCAs at cell fractions less than 3% were excluded, the most frequent autosomal mCAs for the AA and EA ancestry groups were observed on chromosome 11, 12, and 20 (**Sup. Figure 3**). Independent of cell fraction, the rate of mCAs on chromosome 13q was higher for the EA group compared to the AA group (P=0.04) (**Figure 1B)**. Of note, chromosome alterations on 13q are associated with chronic lymphocytic leukemia (CLL) which has a higher incidence in EA compared to other ancestry groups.^17,18^ Across genetic ancestries, the majority of mCAs on chromosomes 1, 11, 14, and 20 were CN-LOH or losses, while gains were commonly observed for chromosome 12 (**Figure 1B**), another signature of CLL.

### Germline predictors of chromosomal alterations across the genome

Next, we performed a WGS-based genome-wide association analysis (GWAS) between germ-line variants observed in TOPMed and presence of an mCA, separately for autosomal (N= 67,518) and chrX (N=41,864) mCA (see **Methods**). For each sample, we defined the phenotype as presence/absence of one or more autosomal mCAs and tested against all variants with minor allele count (MAC) >= 5 that passed the quality filters (**Table S5 & Table S6**). Of the 30 variants reported to be associated with presence of an autosomal mCA in Loh et al.,^6^ we replicated 8 associations in the *TERT* gene locus (**Table S6**) at a nominal significance level. We also ran a GWAS with presence of an mCA on chrX in females only. No single variant was significant at a genome-wide Bonferroni corrected p-value threshold (5e-09) for the GWAS of autosomal mCAs or chrX mCAs in females.

Next, we performed *cis*-analyses (see **Methods**), testing for association between presence of an mCA and germline variants (with MAC >= 5) on the same chromosome arm as the mCA (**Figure 3A**). Based on frequency in our dataset and importance from the literature, we defined the following mCA binary phenotypes: CN-LOH at *MPL*, CN-LOH at *ATM*, 11q CN-LOH, 1p CN-LOH, 12p gain, 12q gain, 14q CN-LOH, and 20q loss (**Table S5**). We found a 3-prime UTR variant at the *ATM* gene (rs3092836) that was significantly associated with mCA at *ATM* (OR=26.41, p=0.0013) and multiple other variants in *ATM* and *MPL* that were associated at a nominal (p<0.05) level (**Table S7**). To determine whether these variants were selectively located on the haplotype that was duplicated in the CN-LOH events at *MPL* and *ATM*, we conducted an allelic shift analysis as in Loh et al. In contrast to Loh et al., we did not find evidence for an allelic shift at these two loci (**Table S8**). The rs3092836 variant was present at an estimated minor allele frequency (MAF) of 8% in AA, 2% in EAS and HA, and 0.06% in EA. Of the variants with a nominal association with CN-LOH at *ATM* and *MPL*, several varied by ancestry with some having MAF greater than 5% in AA but less than 0.1% in EA. Some variants present at MAC < 20 were estimated to have a large effect in *cis*-association analyses of CN-LOH at *ATM* and *MPL*; however, determining significance posed some challenges due to the limited number of individuals with these rare variants (**Table S9**). One of these variants included rs56009889 at the *ATM* locus (OR = 92, P = 2.5e-8, MAC = 7). Another rare variant included a splice donor variant (rs146249964) that was previously reported to be associated with CN-LOH of *MPL*.^6^ Our analysis replicated the association (OR = 296, P = 9.5e-8, MAC = 7) supporting the role of this variant in clonal expansions. Further evidence of its role in hematopoiesis come from clinical reports of this variant as pathogenic/likely pathogenic in individuals with congenital amegakaryocytic thrombocytopenia^19,20^ particularly among the Ashkenazi Jewish population.

**Figure 3.**
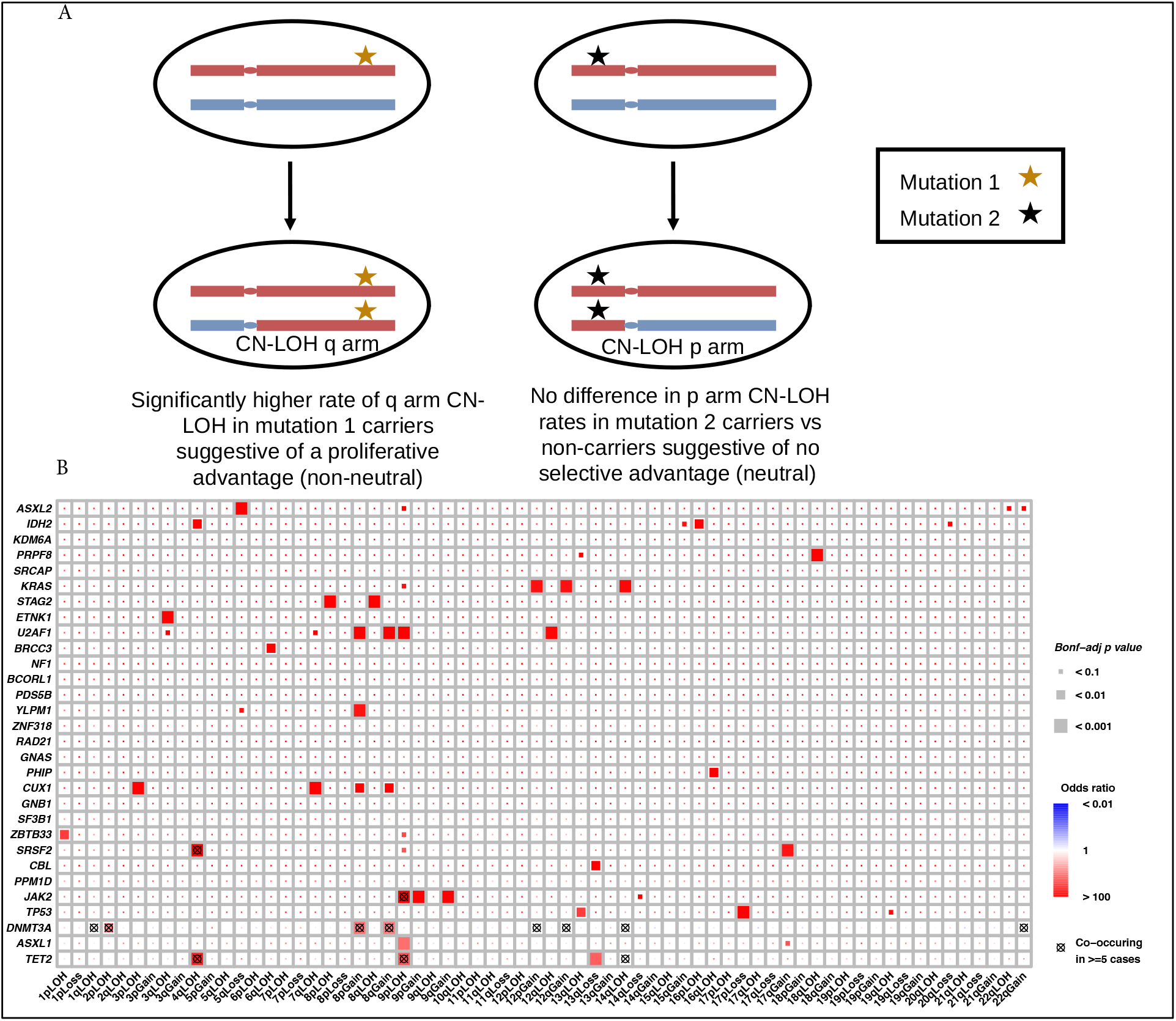
Co-occurrence of mCA and CHIP mutations. **A)** Schematic of how a CHIP mutation may coincide with a CN-LOH event, leading to a proliferative advantage (left), or no selective advantage (right). **B)** Co-occurrence of CHIP and mCA mutations in 30 CHIP genes and 67 mCA events.

To further investigate rare germline variants for association with autosomal and chromosome X mCAs, we implemented gene-centric aggregate rare variants tests for all variants with MAF<1% (see **Methods**). We found 18 statistically significant associations between a burden of rare-variants and presence of mCAs (**Table 1, Figure S9**). The majority of the associations were driven by variants in *ATM* or *MPL*. Of these 18 associations, 11 were at or near *ATM* or *MPL* and were associated with CN-LOH at *MPL* (or 1p CN-LOH) or with CN-LOH at *ATM* (or 11q CN-LOH event). The remaining signals were located at *DCPS* (with any autosomal mCA), *ADAM17* (with any autosomal mCA), *PPP1R16B* (with any autosomal mCA), *TET2* (with any autosomal mCA), and *OR4C16* (with chrX mCA).

**Table 1:** Associations between burden of rare-variants and mCAs.

| Outcome | Variant grouping strategy | chr | Ensembl Gene ID | Gene | Number of variants | MAC | Odds ratio [95% CI] | p | Significance threshold |
| --- | --- | --- | --- | --- | --- | --- | --- | --- | --- |
| autosomal mCA | coding_filter1 | 1 | ENSG00000117400.17 | MPL | 71 | 505 | 1.05 [1.03 - 1.07] | 1.40E-07 | 3.10E-06 |
| autosomal mCA | coding_filter1 | 11 | ENSG00000110063.9 | DCPS | 80 | 626 | 1.04 [1.03 - 1.06] | 4.80E-07 | 3.10E-06 |
| autosomal mCA | coding_filter1 | 2 | ENSG00000151694.13 | ADAM17 | 33 | 72 | 1.12 [1.07 - 1.17] | 1.00E-06 | 3.10E-06 |
| autosomal mCA | coding_filter1 | 20 | ENSG00000101445.9 | PPP1R16B | 14 | 27 | 1.22 [1.12 - 1.31] | 9.60E-07 | 3.10E-06 |
| autosomal mCA | coding_filter1 | 4 | ENSG00000168769.13 | TET2 | 163 | 171 | 1.06 [1.04 - 1.09] | 1.50E-08 | 3.10E-06 |
| autosomal mCA | coding_noncoding_filter1 | 1 | ENSG00000117400.17 | MPL | 71 | 505 | 1.05 [1.03 - 1.07] | 1.40E-07 | 1.40E-06 |
| autosomal mCA | coding_noncoding_filter1 | 11 | ENSG00000110063.9 | DCPS | 82 | 630 | 1.04 [1.03 - 1.06] | 6.30E-07 | 1.40E-06 |
| autosomal mCA | coding_noncoding_filter1 | 20 | ENSG00000101445.9 | PPP1R16B | 14 | 27 | 1.22 [1.12 - 1.31] | 9.60E-07 | 1.40E-06 |
| chr X mCA | coding_filter1 | 11 | ENSG00000279514.2 | OR4C16 | 21 | 174 | 1.04 [1.03 - 1.06] | 2.40E-06 | 3.50E-06 |
| 11q CN-LOH | coding_filter1 | 11 | ENSG00000149311.18 | ATM | 23 | 30 | 1.17 [1.12 - 1.23] | 9.90E-12 | 8.50E-04 |
| 11q CN-LOH | coding_noncoding_filter1 | 11 | ENSG00000149311.18 | ATM | 23 | 30 | 1.17 [1.12 - 1.23] | 9.90E-12 | 9.70E-05 |
| 11q CN-LOH | coding_noncoding_filter1 | 11 | ENSG00000254909.1 | AP003392.2 | 62 | 127 | 1.06 [1.03 - 1.09] | 8.30E-05 | 9.70E-05 |
| 1p CN-LOH | coding_filter1 | 1 | ENSG00000117400.17 | MPL | 15 | 30 | 1.33 [1.27 - 1.39] | 1.20E-32 | 5.50E-04 |
| 1p CN-LOH | coding_noncoding_filter1 | 1 | ENSG00000117400.17 | MPL | 15 | 30 | 1.33 [1.27 - 1.39] | 1.20E-32 | 6.90E-05 |
| ATM CN-LOH | coding_filter1 | 11 | ENSG00000149311.18 | ATM | 22 | 29 | 1.17 [1.11 - 1.22] | 1.30E-10 | 5.00E-02 |
| ATM CN-LOH | coding_noncoding_filter1 | 11 | ENSG00000149311.18 | ATM | 22 | 29 | 1.17 [1.11 - 1.22] | 1.30E-10 | 5.00E-02 |
| MPL CN-LOH | coding_filter1 | 1 | ENSG00000117400.17 | MPL | 15 | 28 | 1.33 [1.26 - 1.39] | 5.40E-31 | 5.00E-02 |
| MPL CN-LOH | coding_noncoding_filter1 | 1 | ENSG00000117400.17 | MPL | 15 | 28 | 1.33 [1.26 - 1.39] | 5.40E-31 | 5.00E-02 |

### Co-occurence of mCAs and CHIP somatic mutations

We interrogated the link between large structural alterations and single nucleotide mutations by tracking the co-occurence of somatic mutations in known CHIP genes that were called in Bick et al. 2020 (N = 3823) and mCAs (N=8402). Overall, individuals with CHIP were more likely to also carry an autosomal mCA (OR=2.76) or an mCA on chromosome X (OR=1.38, **Table S10**). We observed “two-hits” at a number of cancer associated genes (**Figure 3**). These included CHIP mutations co-occurring with CN-LOH at *TET2* (4q), *DNMT3A* (2p), *JAK2* (9p) and *CUX1* (7q). For *TP53*, we observe significant co-occurrence of somatic mutations with loss of chromosome arm 17p where *TP53* is located. Additionally, we observe significant co-occurence of somatic mutations in the *SRSF2* and *KRAS* with gains of these genes. We make a similar observation for the *IDH2* oncogene (15q gain and *IDH2* somatic mutation) however it is marginally significant. Loss of 13q, a common CLL chromosomal alteration, showed significant co-occurrence with somatic mutations in *TET2* and *CBL*.^*17*^ Chromosome 8 gains displayed significant co-occurrence with somatic mutations in *DNMT3A, CUX1*, and *U2AF1*.

### Association between mCAs and hematologic traits and cancers

As has been previously reported, presence of autosomal mCAs at high clonal fractions increases the risk of blood cancers by >10-fold.^4^ Given the high sensitivity of our mCA calls, we investigated the association between autosomal mCAs, chrX mCAs, and mCAs at either high (>= 3%) or low clonal fraction(<3%) with both myeloid (52 cases and 7691 controls) and lymphoid (215 cases and 7291 controls) malignancies (**Table 2**). Autosomal mCAs were associated with an increased risk for lymphoid cancers (OR=2.94, p=4.73e-07) with a stronger effect for high clonal fraction (CF) mCAs (OR=3.78, p=1.46e-07). There were no associations between low CF mCAs or chrX mCAs with lymphoid cancers. Similar to the analysis in Niroula et al.,^21^ we found an even stronger association with lymphoid cancers when we only considered autosomal mCAs that were classified as “lymphoid” (OR=5.64, p=6.60e-09). For myeloid cancers, we found stronger associations with autosomal (OR=5.42, p=1.35e-05) and high CF mCAs (OR=7.77, p=1.74e-06) and no associations with low CF or chrX mCAs. To assess the possibility whether these associations were implicating mCAs as biomarkers for early, sub-clinical disease, we re-ran the associations excluding individuals with cytopenias or cytoses (see **Methods**). The associations with lymphoid cancers were significantly attenuated with these exclusions in place, but the associations with myeloid cancers remained (**Sup. Figure 6**). Due to a paucity of “myeloid” mCAs and myeloid cancer cases, we did not test the association between myeloid mCAs and myeloid cancer. We did not detect an association between autosomal or chrX mCAs with risk for myelodysplastic syndrome (**Table 2**).

**Table 2:** Associations between myeloid and lymphoid malignancies and presence/absence of mcAs.

| Disease | Variable | OR | P-value | n-cases | n-controls |
| --- | --- | --- | --- | --- | --- |
| Lymphoid | autosomal mCA | 2.94 | 4.73E-07 | 215 | 7691 |
| Lymphoid | high CF* autosomal mCA | 3.78 | 1.46E-07 | 215 | 7691 |
| Lymphoid | low CF* autosomal mCA | 1.94 | 7.30E-02 | 215 | 7691 |
| Lymphoid | chrX mCA | 1.23 | 2.96E-01 | 215 | 7691 |
| Lymphoid | autosomal L-mCA | 5.64 | 6.60E-09 | 215 | 7691 |
| Myeloid | autosomal mCA | 5.42 | 1.35E-05 | 52 | 7691 |
| Myeloid | high CF* autosomal mCA | 7.77 | 1.74E-06 | 52 | 7691 |
| Myeloid | low CF* autosomal mCA | 2.61 | 1.92E-01 | 52 | 7691 |
| Myeloid | chrX mCA | 1.08 | 8.56E-01 | 52 | 7691 |
| MDS | autosomal mCA | 1.59 | 1.68E-01 | 112 | 9828 |
| MDS | chrX mCA | 1.11 | 6.99E-01 | 112 | 9828 |
| *high CF mCA was defined as having estimated clonal fraction $\geq 0.03$. | | | | | |
| *low CF mCA was defined as having estimated clonal fraction $< 0.03$. | | | | | |

To investigate the broader impact that inflammatory and behavioral risk factors for blood cancers may have on mCAs, we ran association tests between mCAs and 19 blood cell traits, body mass index (BMI), C-reactive protein (CRP) levels, Interleukin-6 (IL6) levels, and smoking status (see **Methods**) in up to 49,353 individuals. Of the 19 blood cell traits, we observed significant associations between mCA carrier status and levels of lymphocytes, neutrophils, and total white cells (**Table 3**). Specifically, autosomal mCAs were associated with an increase in both lymphocyte counts (beta=0.025) and total white cell counts (beta=0.018); the associations were stronger when we considered mCAs at high clonal fraction only (beta=0.027 for total white cell counts, beta=0.036 for lymphocyte counts). ChrX mCA status was associated with a *decrease* in neutrophil counts (beta = −0.021) and percentages (beta = −0.025), even after adjusting for potential confounding by the Duffy null variant, but was associated with an *increase* in lymphocyte counts (beta = 0.030) and percentages (beta = 0.022) (**Table 3**). We did not observe statistically significant associations between autosomal or chrX mCAs with smoking status, levels of CRP, or levels of IL6 (**Table S11**). Although we did not see a statistically significant association between autosomal mCAs and smoking, the effect estimate (OR=1.67) was similar to that from a previous study.^22^ Finally, we observe d that the presence of autosomal mCAs was associated with a significant decrease in BMI (beta = −0.015, p-value = 0.002). Note that, given the data on hand, we were not able to determine the causal direction of the effects, i.e., we cannot say whether the mCA preceded the risk factor or if the risk factor preceded the mCA.

**Table 3:**
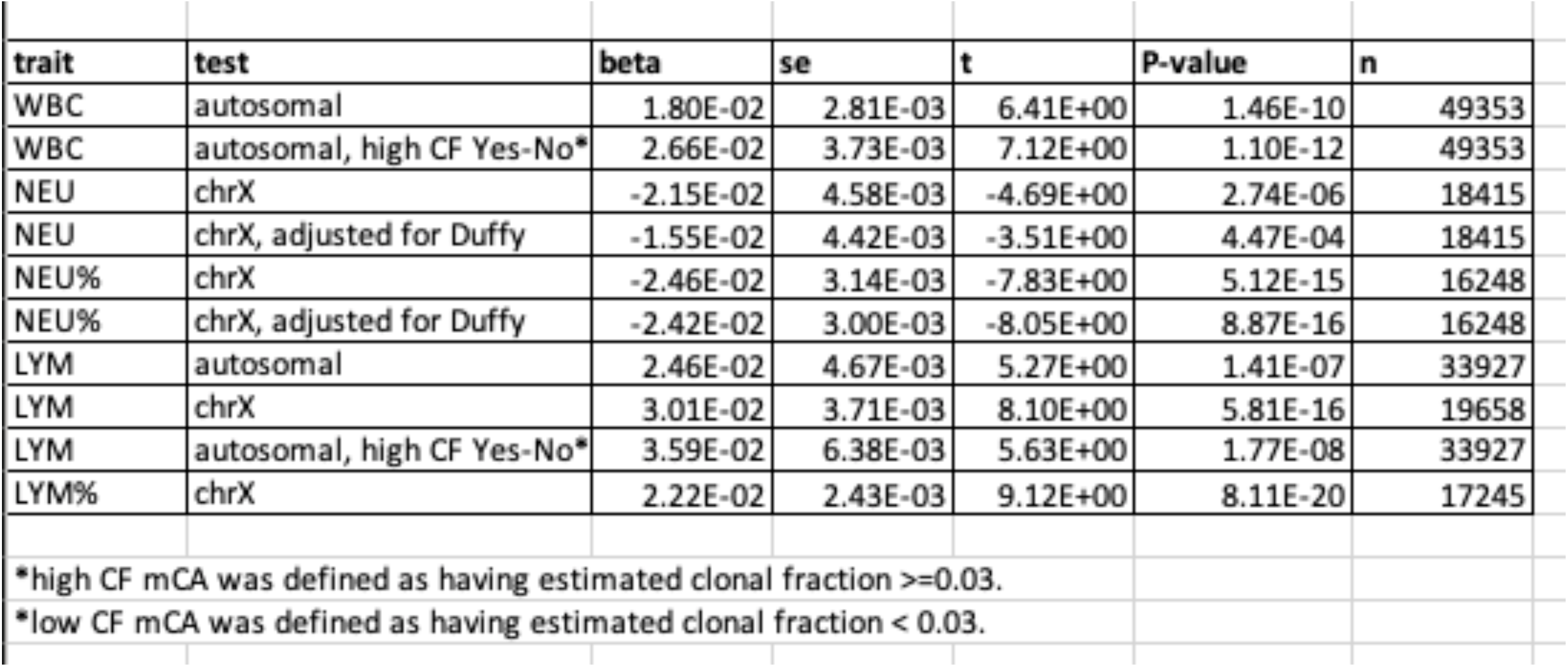
Associations between quantitative blood cell counts and mCA phenotypes.

## DISCUSSION

In this study, we profiled the mCA landscape across an ancestrally diverse set of samples with WGS data to investigate the genomic distribution of mCAs and their germline genetic drivers. Additionally, we detailed the associations of mCAs with incident hematologic cancer, blood cell counts, and markers of inflammation. We observed differences in rates of mCAs across ancestry groups, confirming previous reports of higher prevalence of autosomal mCAs in individuals of EA ancestry relative to AA and EAS ancestry populations.^16,23^ We found a lower autosomal mCA rate in HA ancestry individuals compared to EA. For the first time, we showed that both AA and HA populations have higher rates of chrX mCAs compared to EA. Importantly, these cross-ancestry comparisons were confirmed via a robust down-sampling procedure that removed potential confounding due to differential rates of heterozygosity across ancestries.

The incidence of leukemia is higher in males compared to females, a difference observed across race/ethnicity groups (non-Hispanic White, non-Hispanic Black, Hispanic, and non-Hispanic Asian /Pacific Islander) https://seer.cancer.gov (2019 age-adjusted incidence). Within these race/ethnicity groups, rates in Whites are the highest with a rate of 21 (per 100,000) in males and 13 in females. These rates are lower in Hispanics and Blacks (14 for males and 10 for females). The lowest rates are observed for non-Hispanic Asians (11 for males and 7 for females). Although genetic ancestry is not interchangeable with race/ethnicity, they are correlated. In this multi-ancestry panel from TOPMed we observed that autosomal mCAs rates across ancestry groups follow similar patterns observed for the incidence of leukemia across race/ethnicity groups. This finding supports the use of autosomal mCAs as an intermediate phenotype to study environmental and genetic drivers of blood cancer and as a biomarker for risk.

Both our study and previous studies of European and Japanese populations have uncovered germline variants that increase risk of mCAs, both in-*cis* and in-*trans*. In our cohort, we replicated previous associations at the SNP- and gene-level, demonstrating that rare variants from HA and AA populations are also associated with mCAs. Although some of these variants are ancestry specific, they share molecular drivers, for example with rare variants of large effects driving *cis*-association of mCAs spanning *MPL* and *ATM*. These results motivate further investigation to determine if the observation of an association of a germline variant with mCAs would lend support for classification of a variant as pathogenic in patients with blood cancer, particularly in populations that are under-represented in genetic variant databases.

Relative to what is observed in EA, rates of heterozygosity are higher in the HA and AA admixed populations.^24^ Although this difference presents an advantage for detection of mCAs at lower cell fractions in AA and HA admixed populations, we demonstrate the importance of taking this into account when comparing mCA rates across individuals of different ancestries. From our analyses, this difference in sensitivity was most impactful in detection of chrX mCAs, which, relative to autosomal mCAs, were present at lower cell fractions. The high rates of chrX mCAs (particularly chrX losses) but at overall lower cell fractions suggests that clones with chrX mCAs may arise due to relatively weak positive selection of clones with these mutations and/or possibly high rates of chrX mis-segregation during cell divisions. Both of these explanations are supported by recent work which has shown a significant decrease in hematopoietic clonal diversity in elderly individuals (>75 years) and clonal expansions detectable through single cell sequencing starting before the age of 40.^25^ The observation of chrX mCAs at overall lower cell fractions and higher rates than autosomal mCAs suggests that autosomal mCAs may be under stronger positive selection relative to chrX mCAs which is further supported by overall higher rates of chrX mCAs in AA and HA relative to EA although the incidence of leukemia is higher in EA.

Our study was the first of its kind to implement haplotype-based mCA detection methods on large-scale WGS data from a population-based cohort. With recent enhancement to the MoChA mCA calling software, we were able to detect mCAs with much lower clonal fractions compared to array-based datasets. Our pipeline (https://github.com/auerlab/TOPMed-mCA) relied on substantial post-hoc filtering of mCA calls. In particular, we discarded many small mCA calls due to our inability to distinguish gains, loss, and CN-LOH events for mCAs < 1Mb in size. This decrease in sensitivity for small mCAs may be possible to overcome with higher coverage and or improvements in the detection methods. The WGS data provided a significant advantage over SNP arrays in permitting simultaneous investigation of mCAs, germline variants (including rare/private variants), and CHIP.

This effort represents the first large-scale effort to understand the co-occurrence of distinct forms of mosaicism in individuals of diverse ancestries. Prior work characterizing CHIP and mCAs has focused on individuals of East Asian ancestry, white British individuals or among individuals with solid tumors. Similar to these efforts, we found that CN-LOH co-occurring with CHIP mutations is a common mechanism through which *TET2* (4q), *DNMT3A* (2p), *JAK2* (9p), *CUX1* (7q) and *TP53* (17p) acquire a competitive advantage. We also note that loss of 13q, a common alteration in CLL, co-occurred with CHIP mutations in *TET2* and *CBL*, which may explain how CHIP mutations, despite leading to a myeloid bias, may also predispose to lymphoid malignancy through co-occurring mutations. To date, CHIP and mCAs have largely been considered independently. Saiki et al.^26^ reported that individuals with both mCAs and CHIP have worse outcomes. An important area of further investigation will focus on the distinction between heterozygosity and homozygosity at a CHIP locus and disease consequences. It is tempting to speculate that patients with clones that make up the same fraction of blood that are homozygous for CHIP mutations due to concomitant CN-LOH may have worse prognosis than individuals with a single mutation.

## METHODS

### Study population

We included 67,390 participants from 19 TOPMed studies: Genetics of Cardiometabolic Health in the Amish (Amish, n=1,109)^27^, Atherosclerosis Risk in Communities Study (ARIC, n=3,780)^28^, Barbados Genetics Asthma Study (BAGS, n=980), Mount Sinai BioMe Biobank (BioMe, n=9,392)^29^, Coronary Artery Risk Development in Young Adults (CARDIA, n=3,293)^30^, Cleveland Family Study (CFS, n=1,281), Cardiovascular Health Study (CHS, n=3,517)^31^, Genetic Epidemiology of COPD Study (COPDGene, n=10,050)^32^, Framingham Heart Study (FHS, n=4,007)^33^, Genetic Studies of Atherosclerosis Risk (GeneSTAR, n=1,733)^34^, Genetic Epidemiology Network of Arteriopathy (GENOA, n=1,157), Genetics of Lipid Lowering Drugs and Diet Network (GOLDN, n=942), Hispanic Community Health Study - Study of Latinos (HCHS_SOL, n=3,857)^35^, Hypertension Genetic Epidemiology Network (HyperGEN, n=1,865), Jackson Heart Study (JHS, n=3,317)^36^, Multi-Ethnic Study of Atherosclerosis (MESA, n=5,222)^37^, Vanderbilt BioVU study of African Americans (VU_AF, n=1,085), Women’s Genome Health Study (WGHS, n=108), and Women’s Health Initiative (WHI, n=10,695)^38^. The 67,390 TOPMed participants were categorized into discrete ancestry subgroups using the HARE (harmonized ancestry and race/ethnicity) machine learning algorithm,^39^ which uses genetically inferred ancestry to refine self-identified race/ethnicity and impute missing racial/ethnic values. The ancestry composition in this study was 57% European, 30% African, 11% Hispanic/Latino, and 2% Asian (**see Table S12**). Further descriptions of the design of the participating TOPMed cohorts and the sampling of individuals within each cohort for TOPMed WGS are provided in the section “Participating studies” under **Supplemental Methods**. All studies were approved by the appropriate institutional review boards (IRBs) and informed consent was obtained from all participants.

### WGS data

WGS was performed as part of the NHLBI TOPMed program. The WGS was performed at an average depth of 38X by six sequencing centers (Broad Genomics, Northwest Genome Institute, Illumina, New York Genome Center, Baylor, and McDonnell Genome Institute) using Illumina X10 technology and DNA from blood. Here we report analyses from ‘Freeze 8,’ for which reads were aligned to human-genome build GRCh38 using a common pipeline across all centers. To perform variant quality control (QC), a support vector machine (SVM) classifier was trained on known variant sites (positive labels) and Mendelian inconsistent variants (negative labels). Further variant filtering was done for variants with excess heterozygosity and Mendelian discordance. Sample QC measures included: concordance between annotated and inferred genetic sex, concordance between prior array genotype data and TOPMed WGS data, and pedigree checks. Additional details can be found in Taliun et al.^24^

### Detection of mCAs

Detection of mCAs was performed on the WGS-based genotype and read depth data. The mCA call set was generated using the MOsaic CHromosomal Alterations (MoChA v1.11) caller. This approach utilizes phased genotypes, coverage, and B allele frequency (BAF) at heterozygous sites for detection of mCAs. Input data at heterozygous markers came from a previous analysis of the TOPMed cohort as outlined in Taliun et al.^24^ However, not all variants were included in the analyses. First, heterozygous markers with a minor allele frequency less than 1% and those where the read depth of either allele was less than 5 were removed. Second, we removed markers within germline copy number variants (CNVs) previously generated in TOPMed. Third, when more than 1 marker was present in a 1,000 base pair genomic region, then only one marker was retained. The MoChA caller was run with the extra option “--LRR-weight 0.0 --bdev-LRR-BAF 6.0” to disable the LRR+BAF model. The resulting mCA calls were filtered by excluding 1) those that span less than 2000 informative markers i.e. heterozygous sites; 2) those with lod score less than 5, 3) those on chromosome X but with inferred sex “unknown”, 4) those with estimated relative coverage higher than 2.9, 5) those with BAF deviation larger than 0.16 and relative coverage higher than 2.5. Steps 4 and 5 are used to exclude putative germline duplications. Classification of mCAs as lymphoid or myeloid was performed following criterion from Niroula et al.^21^

### Down-sampling

The total number of heterozygous sites can affect the power for detection of mCAs as the mCA calling method relies on heterozygous sites for detecting imbalances in the parental haplotypes.^5,40^ The AA, HA, and EA groups had on average higher number of heterozygous sites compared to the EAS group (**Table S4**); therefore to adjust for this difference we down-sampled heterozygous sites in the AA, HA, and EA groups and then used those data to generate mCA calls and re-asses reported associations of mCAs with ancestry. The down-sampling was conducted by matching the distribution of heterozygous sites for AA, HA, and EA groups to that of the EAS group. This adjustment was done separately for females and males. For example, if a HA female sample had 925,935 heterozygous markers which is equivalent to the 50th percentile for HA females, then heterozygous markers were removed at random across the genome until the sample had 749,959 markers which is equal to the 50th percentile for EAS females.

### Comparisons of mCAs across ancestries

Subsequent to down-sampling, we investigated possible batch effects that may have influenced mCA detection rates across both autosomes and chrX. After adjusting for age, age^2^, sex, and ancestry, a variable representing “study” had no effect on autosomal mCA detection, though we did find a study effect for chrX mCA detection. Therefore, in all of our analyses comparing mCA detection across ancestries, we included age, age^2^, sex, and study as covariates.

### Association Analyses

We performed a WGS-based genome-wide association analysis (GWAS) between germ-line variants observed in TOPMed and presence of an mCA, separately for autosomal (N=67,518) and chrX (N=41,864) mCAs. For each sample, we defined the phenotype as presence/absence of one or more autosomal mCAs and tested against all variants with minor allele count (MAC) >= 5 that passed the quality filters. Samples with uncertain identity or poor quality were excluded from analysis. Principal components and genetic relatedness estimates were calculated using PC-AiR^41^ and PC-Relate^42^, as described previously in Hu et al.^43^ QC replicates or duplicate samples were removed after selecting the sample with the highest average autosomal depth rate. All logistic regression analyses included age, age squared, sex, study, and genetic ancestry as covariates. The final sample set included 5 genetic ancestry categories consisting of African American (AA), European American (EA), East Asian (EAS), Hispanic American (HA), and a group of 1099 samples that were characterized as having “unknown” ancestry. To test for association of sex with specific mCA types, for example 20q loss, we first conducted a chi-square test (R chisq.test, simulate.p.value = TRUE, B = 100,000). For mCAs types with marginal significance (p < 0.1) we then conducted logistic regression to test for association using a Bonferroni correction to account for the 156 independent tests.

We performed genetic association tests in *cis* and in *trans* using a generalized linear mixed model (GLMM) approach using the GMMAT method^44^ as implemented in the GENESIS software.^45^ For each analysis, a null model assuming no association between the outcome and any variant was fit, adjusting for sex, age, study-sequencing phase, and the first 11 PCs to capture genetic ancestry. A 4th degree sparse empirical kinship matrix (KM) computed with PC-Relate was included as a random effect to account for genetic relatedness among participants. The residuals from this null model were then used to perform genome-wide score tests of genetic association.

For the *trans* association analyses, we defined cases as those with a detectable mCA and tested all genetic variants with MAC >= 20 and had less than 10% of samples freeze-wide with sequencing read depth < 10 at that particular variant.

For the *cis* associations (i.e., variants within the same genomic locus as the mCA), we identified 8 genomic loci of interest which included the *ATM* and *MPL* genes as well as chromosome arms with recurrent autosomal mCAs (n > 100) which included 14q CN-LOH, 1q CN-LOH, 11q CN-LOH, 12p gain, 12q gain, 20q loss, 1q CN-LOH. Cases were defined as those with an mCA call spanning the chromosome arm or gene, while controls were defined at those without any mCA calls on the chromosome arm tested. Because the case-control ratio was highly unbalanced for these analyses, we matched cases to controls using study, sequencing phase, sex, and age to obtain a 1:10 ratio before fitting the null model. We tested all variants that passed the quality filters and had MAF >= 0.01. We defined a significance threshold of p < 0.05 / (effective number of variants), where the effective number of variants tested was calculated using simpleM.^46^

For the *ATM* and *MPL* gene analyses, we further filtered variants based on annotations. Variants were annotated using ANNOVAR (v2019-10-24) and selected for use based on their presence in exons, and/or potential involvement in splicing. In addition to canonical splice sites, we also tested variants +/- 6 bp from exon boundaries, as well as less-canonical splice sites identified by SPIDEX.^47^ To specifically account for promoters, we identified promoters for these two genes using the Eukaryotic Promoter Database, and included variants found in the promoter region. Similar to the *cis* association analyses, we defined a significance threshold of p < 0.05 / (effective number of variants), where the effective number of variants tested was calculated using simpleM. We also ran secondary analyses of these variants with a lower MAC threshold (MAC >= 5).

In addition to single variant testing, we conducted gene-based aggregate tests to assess the cumulative effect of rare variants on mCA presence. Variants were aggregated by gene using the GENCODE v29 gene model. We used two strategies for filtering variants. For both strategies, variants were first filtered to MAF < 0.01 in the sample set being tested. The first strategy (coding_filter1) includes only high confidence predicted LoF variants inferred using LOFTEE (https://github.com/konradjk/loftee) and missense variants filtered using MetaSV score > 0.^48^ The second strategy (coding_noncoding_filter1) includes all variants from the first strategy plus additional regulatory variants. Regulatory variants were included if they overlapped with enhancer(s) or promoters linked to a gene using GeneHancer,^49^ or 5 Kb upstream of the Transcription start site. Within these regions only those variants were retained which had Fathmm-XF score > 0.5 or overlap with regions labeled as either “CTCF binding sites” or “Transcription factor binding sites” as annotated by the Ensembl regulatory build annotation.^50^ Results were filtered to only those aggregation units with a cumulative MAC >= 20. We defined the significance threshold as p < 0.05 / (number of aggregation units tested).

The annotation based variant filtering and gene based aggregation was performed using TOPMed freeze 8 WGSA Google BigQuery annotation database on the BiodataCatalyst powered by Seven Bridges platform (http://doi.org/10.5281/zenodo.3822858). The annotation database was built using variant annotations generated by Whole Genome Sequence Annotator (WGSA) version v0.8^51^ and formatted by WGSAParsr version 6.3.8 (<https://github.com/UW-GAC/wgsaparsr>). The GENCODE v29 gene model based varint consequences were obtained from Ensembl Variant effect predictor (VEP)^52^ incorporated within WGSA. When using a deleteriousness prediction score, respective author recommended cut points were used to retain likely deleterious variants.

### Co-Occurence Analysis

Co-occurence between CHIP status and mCA status was analyzed as in previous work.^26^ First, CHIP “carriers” (individuals with an observed acquired mutation) were assigned to different categories by the gene where the mutations were located. Carriers of mCAs were assigned to categories based on the location (chromosome, p-arm and q-arm) and the changes of copy numbers (Gain, Loss, and copy neutral LOH) of the mCA. A CHIP carrier or an mCA carrier may have been assigned to different categories if that individual carried multiple CHIP mutations or multiple mCAs. In our analysis, we required there to be at least 10 carriers in each category, leaving 30 CHIP categories and 66 mCA categories for comparison (**Figure 3**). P-values for co-occurence of CHIP and mCA carrier status were obtained via the Wald test (note that 0.5 was added to all cells in the 2×2 table if zero value(s) exists). A Bonferroni correction was implemented to assess significance.

### Analysis of Hematologic Malignancy Data

Due to a paucity of cancer outcome data in all other cohorts, we restricted our analysis of hematologic malignancies to the WHI. We assigned the available hematologic cancer outcomes in the WHI cohort into categories: lymphoid and myeloid cancers. Patients diagnosed as chronic lymphocytic leukemia, non-Hodgkins lymphoma, or multiple myeloma were assigned to the lymphoid group (n=237). All other patients diagnosed with leukemia were assigned to the myeloid group (n=53). We further excluded patients that were diagnosed as having any cancer before blood draw (i.e., the time at which DNA for mCA calling was collected), which reduced the lymphoid cancer case number to 223 and the myeloid cancer case number to 52.

We ran separate logistic regression models to test the association between mCA carrier status and risk for lymphoid or myeloid cancer. Covariates in our model included CHIP carrier status, the interaction between CHIP carrier status and mCA carrier status, age, ancestry, and smoking. Ancestry groups with fewer than 5 cases were not included in this analysis. To determine whether any associations with myeloid or lymphoid malignancies were implicating mCAs as biomarkers for early, sub-clinical disease, we re-ran these logistic regressions excluding individuals with cytopenias or cytoses. Cytopenia and cytosis cases were defined based on the blood cell counts collected at any of three WHI visits, using the definitions in **Supplementary Table S13**.

### Analysis with inflammatory and blood cell traits

A description of the measurement and quality control of the blood-cell and inflammation traits can be found in Stilp et al.^53^ Each trait was defined as follows: Hematocrit (HCT) is the percentage of volume of blood that is composed of red blood cells. Hemoglobin (HGB) is the mass per volume (grams per deciliter) of hemoglobin in the blood. Mean corpuscular hemoglobin (MCH) is the average mass in picograms of hemoglobin per red blood cell. Mean corpuscular hemoglobin concentration (MCHC) is the average mass concentration (grams per deciliter) of hemoglobin per red blood cell. Mean corpuscular volume (MCV) is the average volume of red blood cells, measured in femtoliters (fL). RBC count is the count of red blood cells in the blood, by number concentration in millions per microliter. Red cell distribution width (RDW) is the measurement of the ratio of variation in width to the mean width of the red blood cell volume distribution curve taken at +/- one CV. Total white blood cell count (WBC), neutrophil (NEU), monocyte (MONO), lymphocyte (LYM), eosinophil (EOS), basophil (BASO) and platelet count (PLT) are defined with respect to cell concentration in blood, measured in thousands/microliter. Because of a typical large point mass at zero, we dichotomized the BASO phenotype at BASO>0. The proportion of neutrophils, monocytes, lymphocytes, or eosinophils were calculated by dividing the respective WBC sub-type count by the total measured WBC. Mean platelet volume (MPV) was measured in fL. C-reactive protein (CRP) was measured in mg/L. Interleukin 6 (IL6) was measured in pg/ML. Body mass index (BMI) was calculated from standing height and weight and smoking was dichotomized as ever/never smoker.

We tested for the association between each of these traits and presence of autosomal mCAs, chrX mCAs, and mCAs at either high (>3%) or low clonal fraction. We ran standard linear models treating each quantitative trait as the dependent variable and presence/absence of mCA as the main independent variable of interest while adjusting for age, sex, ancestry group, and TOPMed study phase. BMI, WBC, NEU, LYM, MONO, and EOS were log transformed. We ran logistic regressions to test the association with smoking and BASO, adjusting for the same set of covariates.

## ACKNOWLEDGEMENTS

Molecular data for the Trans-Omics in Precision Medicine (TOPMed) program was supported by the National Heart, Lung and Blood Institute (NHLBI). The table below presents study specific omics support information. Core support including centralized genomic read mapping and genotype calling, along with variant quality metrics and filtering were provided by the TOPMed Informatics Research Center (3R01HL-117626-02S1; contract HHSN268201800002I). Core support including phenotype harmonization, data management, sample-identity QC, and general program coordination were provided by the TOPMed Data Coordinating Center (R01HL-120393; U01HL-120393; contract HHSN268201800001I).

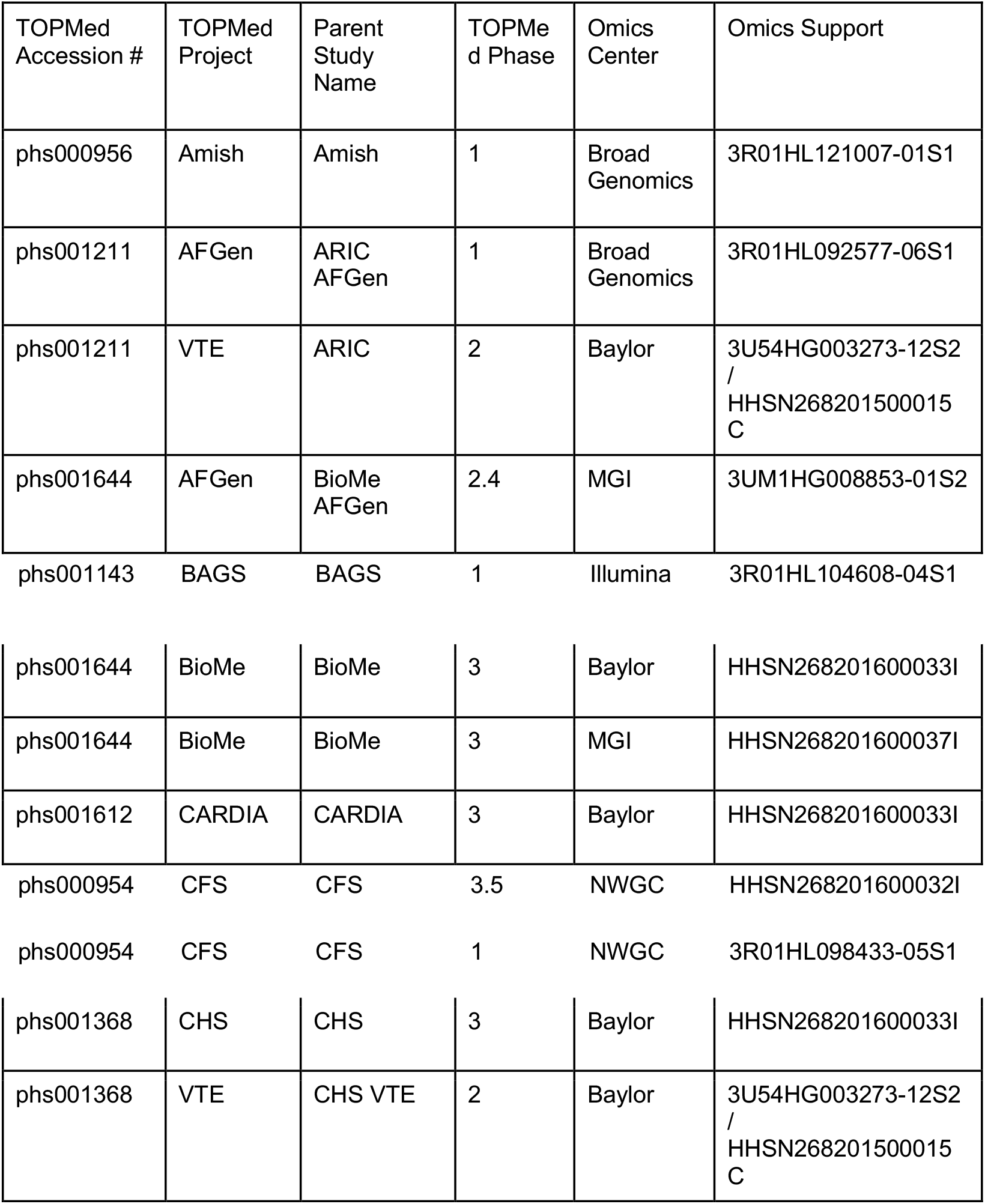
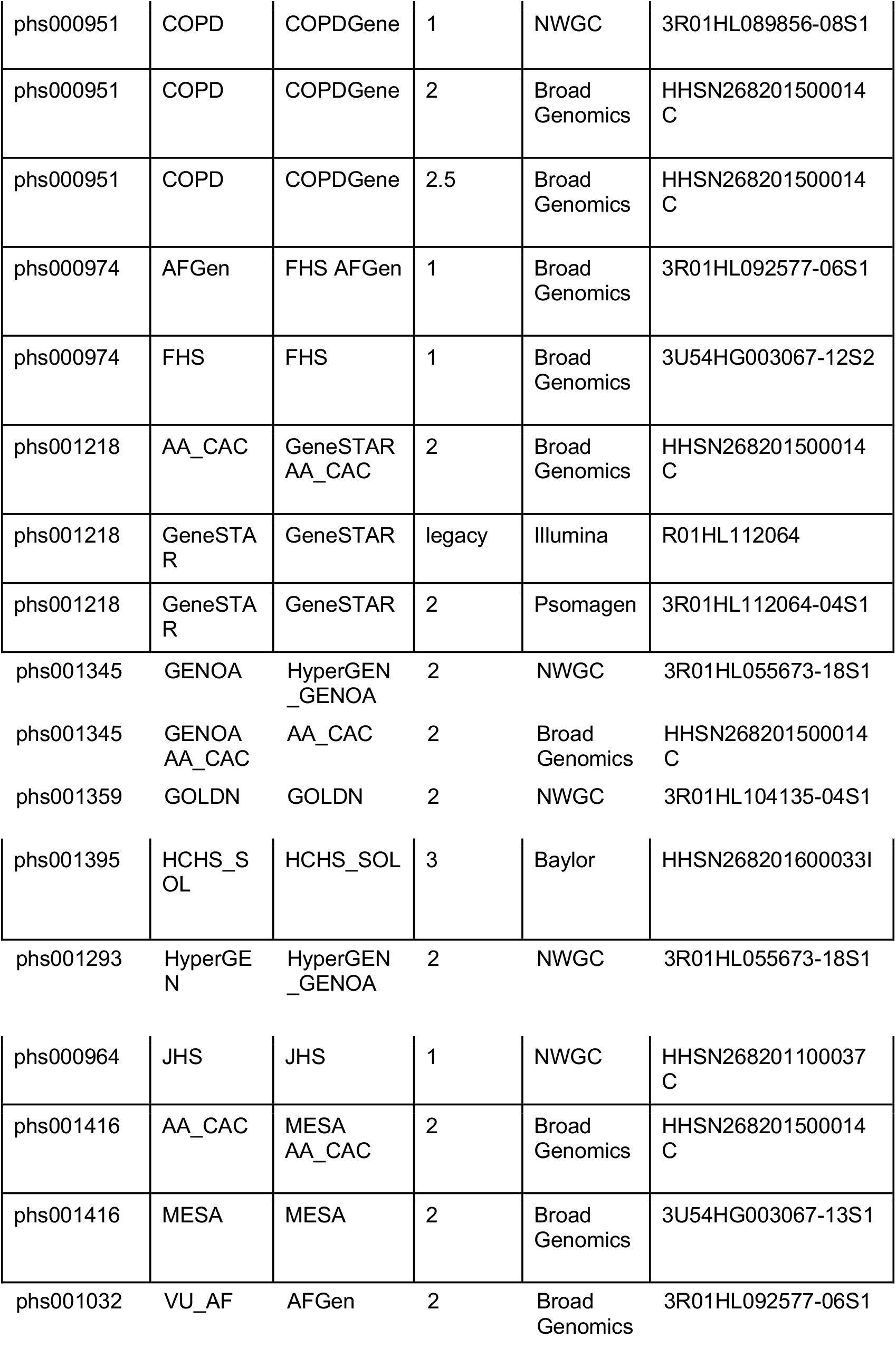
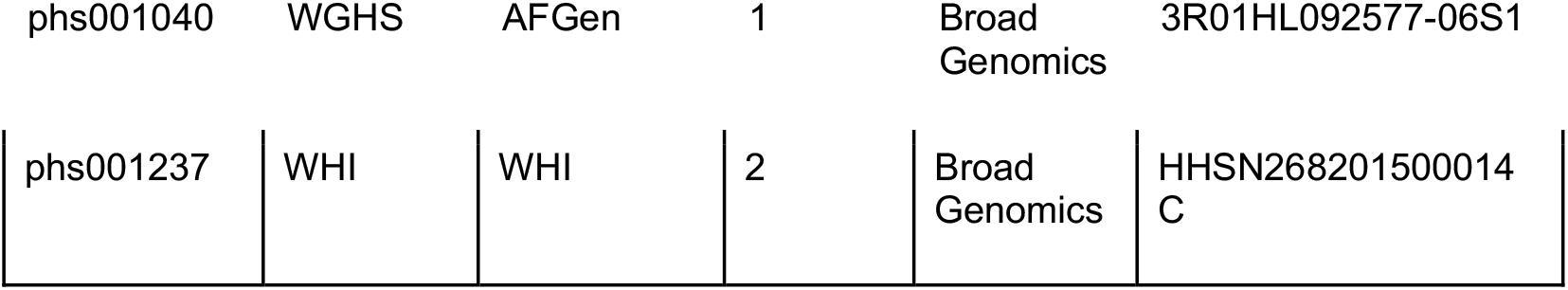

| TOPMed Accession # | TOPMed Project | Parent Study Name | TOPMed Phase | Omics Center | Omics Support |
| --- | --- | --- | --- | --- | --- |
| phs000956 | Amish | Amish | 1 | Broad Genomics | 3R01HL121007-01S1 |
| phs001211 | AFGen | ARIC AFGen | 1 | Broad Genomics | 3R01HL092577-06S1 |
| phs001211 | VTE | ARIC | 2 | Baylor | 3U54HG003273-12S2 / HHSN268201500015C |
| phs001644 | AFGen | BioMe AFGen | 2.4 | MGI | 3UM1HG008853-01S2 |
| phs001143 | BAGS | BAGS | 1 | Illumina | 3R01HL104608-04S1 |
| phs001644 | BioMe | BioMe | 3 | Baylor | HHSN268201600033I |
| phs001644 | BioMe | BioMe | 3 | MGI | HHSN268201600037I |
| phs001612 | CARDIA | CARDIA | 3 | Baylor | HHSN268201600033I |
| phs000954 | CFS | CFS | 3.5 | NWGC | HHSN268201600032I |
| phs000954 | CFS | CFS | 1 | NWGC | 3R01HL098433-05S1 |
| phs001368 | CHS | CHS | 3 | Baylor | HHSN268201600033I |
| phs001368 | VTE | CHS VTE | 2 | Baylor | 3U54HG003273-12S2 / HHSN268201500015C |
| phs000951 | COPD | COPDGene | 1 | NWGC | 3R01HL089856-08S1 |
| phs000951 | COPD | COPDGene | 2 | Broad Genomics | HHSN268201500014C |
| phs000951 | COPD | COPDGene | 2.5 | Broad Genomics | HHSN268201500014C |
| phs000974 | AFGen | FHS AFGen | 1 | Broad Genomics | 3R01HL092577-06S1 |
| phs000974 | FHS | FHS | 1 | Broad Genomics | 3U54HG003067-12S2 |
| phs001218 | AA_CAC | GeneSTAR AA_CAC | 2 | Broad Genomics | HHSN268201500014C |
| phs001218 | GeneSTAR | GeneSTAR | legacy | Illumina | R01HL112064 |
| phs001218 | GeneSTAR | GeneSTAR | 2 | Psomagen | 3R01HL112064-04S1 |
| phs001395 | HCHS_SOL | HCHS_SOL | 3 | Baylor | HHSN268201600033I |
| phs001293 HyperGEN HyperGEN\_GENOA 2 NWGC 3R01HL055673-18S1 |  |  |  |  |  |
| phs000964 | JHS | JHS | 1 | NWGC | HHSN268201100037C |
| phs001416 | AA_CAC | MESA AA_CAC | 2 | Broad Genomics | HHSN268201500014C |
| phs001416 | MESA | MESA | 2 | Broad Genomics | 3U54HG003067-13S1 |
| phs001032 VU\_AF AFGen 2 Broad Genomics 3R01HL092577-06S1 |  |  |  |  |  |
| phs001040 | WGHS | AFGen | 1 | Broad Genomics | 3R01HL092577-06S1 |
| phs001237 | WHI | WHI | 2 | Broad Genomics | HHSN268201500014 C |

Amish: The TOPMed component of the Amish Research Program was supported by NIH grants R01 HL121007, U01 HL072515, and R01 AG18728.

ARIC: The Atherosclerosis Risk in Communities study has been funded in whole or in part with Federal funds from the National Heart, Lung, and Blood Institute, National Institutes of Health, Department of Health and Human Services (contract numbers HHSN268201700001I, HHSN268201700002I, HHSN268201700003I, HHSN268201700004I and HHSN268201700005I). The authors thank the staff and participants of the ARIC study for their important contributions.

BAGS: We gratefully acknowledge the contributions of Pissamai and Trevor Maul, Paul Levett, Anselm Hennis, P. Michele Lashley, Raana Naidu, Malcolm Howitt and Timothy Roach, and the numerous health care providers, and community clinics and co-investigators who assisted in the phenotyping and collection of DNA samples, and the families and patients for generously donating DNA samples to the Barbados Asthma Genetics Study (BAGS). Funding for BAGS was provided by National Institutes of Health (NIH) R01HL104608, R01HL087699, and HL104608 S1.

BioMe: The Mount Sinai BioMe Biobank has been supported by The Andrea and Charles Bronfman Philanthropies and in part by Federal funds from the NHLBI and NHGRI (U01HG00638001; U01HG007417; X01HL134588). We thank all participants in the Mount Sinai Biobank. We also thank all our recruiters who have assisted and continue to assist in data collection and management and are grateful for the computational resources and staff expertise provided by Scientific Computing at the Icahn School of Medicine at Mount Sinai.

CARDIA: The Coronary Artery Risk Development in Young Adults Study (CARDIA) is conducted and supported by the National Heart, Lung, and Blood Institute (NHLBI) in collaboration with the University of Alabama at Birmingham (HHSN268201800005I & HHSN268201800007I), Northwestern University (HHSN268201800003I), University of Minnesota (HHSN268201800006I), and Kaiser Foundation Research Institute (HHSN268201800004I). CARDIA was also partially supported by the Intramural Research Program of the National Institute on Aging (NIA) and an intra-agency agreement between NIA and NHLBI (AG0005).

CFS: The Cleveland Family Study has been supported in part by National Institutes of Health grants [R01-HL046380, KL2-RR024990, R35-HL135818, and R01-HL113338].

CHS: Cardiovascular Health Study: This research was supported by contracts 75N92021D00006, HHSN268201200036C, HHSN268200800007C, HHSN268201800001C, N01HC55222, N01HC85079, N01HC85080, N01HC85081, N01HC85082, N01HC85083, N01HC85086, and grants U01HL080295 and U01HL130114 from the National Heart, Lung, and Blood Institute (NHLBI), with additional contribution from the National Institute of Neurological Disorders and Stroke (NINDS). Additional support was provided by R01AG023629 from the National Institute on Aging (NIA). A full list of principal CHS investigators and institutions can be found at CHS-NHLBI.org. The content is solely the responsibility of the authors and does not necessarily represent the official views of the National Institutes of Health.

COPDGene: The COPDGene project described was supported by Award Number U01 HL089897 and Award Number U01 HL089856 from the National Heart, Lung, and Blood Institute. The content is solely the responsibility of the authors and does not necessarily represent the official views of the National Heart, Lung, and Blood Institute or the National Institutes of Health. COPDGene is also supported by the COPD Foundation through contributions made to an Industry Advisory Board that has included AstraZeneca, Bayer Pharmaceuticals, Boehringer-Ingelheim, Genentech, GlaxoSmithKline, Novartis, Pfizer, and Sunovion. A full listing of COPDGene investigators can be found at: http://www.copdgene.org/directory

FHS: The Framingham Heart Study (FHS) acknowledges the support of contracts NO1-HC-25195 and HHSN268201500001I from the National Heart, Lung and Blood Institute and grant supplement R01 HL092577-06S1 for this research. We also acknowledge the dedication of the FHS study participants without whom this research would not be possible.

GeneSTAR: GeneSTAR was supported by the National Institutes of Health/National Heart, Lung, and Blood Institute (U01 HL72518, HL087698, HL112064, HL11006, HL118356) and by a grant from the National Institutes of Health/National Center for Research Resources (M01-RR000052) to the Johns Hopkins General Clinical Research Center. We would like to thank our participants and staff for their valuable contributions.

GENOA: Support for GENOA was provided by the National Heart, Lung and Blood Institute (U01 HL054457, U01 HL054464, U01 HL054481, R01 HL119443, and R01 HL087660) of the National Institutes of Health.

GOLDN: GOLDN biospecimens, baseline phenotype data, and intervention phenotype data were collected with funding from National Heart, Lung and Blood Institute (NHLBI) grant U01 HL072524. Whole-genome sequencing in GOLDN was funded by NHLBI grant R01 HL104135 and supplement R01 HL104135-04S1.

HCHS/SOL: The Hispanic Community Health Study/Study of Latinos is a collaborative study supported by contracts from the National Heart, Lung, and Blood Institute (NHLBI) to the University of North Carolina (HHSN268201300001I / N01-HC-65233), University of Miami (HHSN268201300004I / N01-HC-65234), Albert Einstein College of Medicine (HHSN268201300002I / N01-HC-65235), University of Illinois at Chicago – HHSN268201300003I / N01-HC-65236 Northwestern Univ), and San Diego State University (HHSN268201300005I / N01-HC-65237). The following Institutes/Centers/Offices have contributed to the HCHS/SOL through a transfer of funds to the NHLBI: National Institute on Minority Health and Health Disparities, National Institute on Deafness and Other Communication Disorders, National Institute of Dental and Craniofacial Research, National Institute of Diabetes and Digestive and Kidney Diseases, National Institute of Neurological Disorders and Stroke, NIH Institution-Office of Dietary Supplements.

HyperGEN: The HyperGEN Study is part of the National Heart, Lung, and Blood Institute (NHLBI) Family Blood Pressure Program; collection of the data represented here was supported by grants U01 HL054472 (MN Lab), U01 HL054473 (DCC), U01 HL054495 (AL FC), and U01 HL054509 (NC FC). The HyperGEN: Genetics of Left Ventricular Hypertrophy Study was supported by NHLBI grant R01 HL055673 with whole-genome sequencing made possible by supplement −18S1.

JHS: The Jackson Heart Study (JHS) is supported and conducted in collaboration with Jackson State University (HHSN268201800013I), Tougaloo College (HHSN268201800014I), the Mississippi State Department of Health (HHSN268201800015I) and the University of Mississippi Medical Center (HHSN268201800010I, HHSN268201800011I and HHSN268201800012I) contracts from the National Heart, Lung, and Blood Institute (NHLBI) and the National Institute on Minority Health and Health Disparities (NIMHD). The authors also wish to thank the staffs and participants of the JHS.

MESA: Whole genome sequencing (WGS) for the Trans-Omics in Precision Medicine (TOPMed) program was supported by the National Heart, Lung and Blood Institute (NHLBI). WGS for “NHLBI TOPMed: Multi-Ethnic Study of Atherosclerosis (MESA)” (phs001416.v1.p1) was performed at the Broad Institute of MIT and Harvard (3U54HG003067-13S1). Centralized read mapping and genotype calling, along with variant quality metrics and filtering were provided by the TOPMed Informatics Research Center (3R01HL-117626-02S1). Phenotype harmonization, data management, sample-identity QC, and general study coordination, were provided by the TOPMed Data Coordinating Center (3R01HL-120393-02S1), and TOPMed MESA Multi-Omics (HHSN2682015000031/HSN26800004). The MESA projects are conducted and supported by the National Heart, Lung, and Blood Institute (NHLBI) in collaboration with MESA investigators. Support for the Multi-Ethnic Study of Atherosclerosis (MESA) projects are conducted and supported by the National Heart, Lung, and Blood Institute (NHLBI) in collaboration with MESA investigators. Support for MESA is provided by contracts 75N92020D00001, HHSN268201500003I, N01-HC-95159, 75N92020D00005, N01-HC-95160, 75N92020D00002, N01-HC-95161, 75N92020D00003, N01-HC-95162, 75N92020D00006, N01-HC-95163, 75N92020D00004, N01-HC-95164, 75N92020D00007, N01-HC-95165, N01-HC-95166, N01-HC-95167, N01-HC-95168, N01-HC-95169, UL1-TR-000040, UL1-TR-001079, UL1-TR-001420, UL1TR001881, DK063491, and R01HL105756. The authors thank the other investigators, the staff, and the participants of the MESA study for their valuable contributions. A fill list of participating MESA investigators and institutes can be found at http://www.mesa-nhlbi.org.

WGHS: The WGHS is supported by the National Heart, Lung, and Blood Institute (HL043851 and HL080467) and the National Cancer Institute (CA047988 and UM1CA182913). The most recent cardiovascular endpoints were supported by ARRA funding HL099355.

WHI: The WHI program is funded by the National Heart, Lung, and Blood Institute, National Institutes of Health, U.S. Department of Health and Human Services through contracts HHSN268201600018C, HHSN268201600001C, HHSN268201600002C, HHSN268201600003C, and HHSN268201600004C.

The views expressed in this manuscript are those of the authors and do not necessarily represent the views of the National Heart, Lung, and Blood Institute; the National Institutes of Health; or the U.S. Department of Health and Human Services.

