## Supplementary material for "Mosaic chromosomal alterations in blood across ancestries via whole-genome sequencing": Figure S9

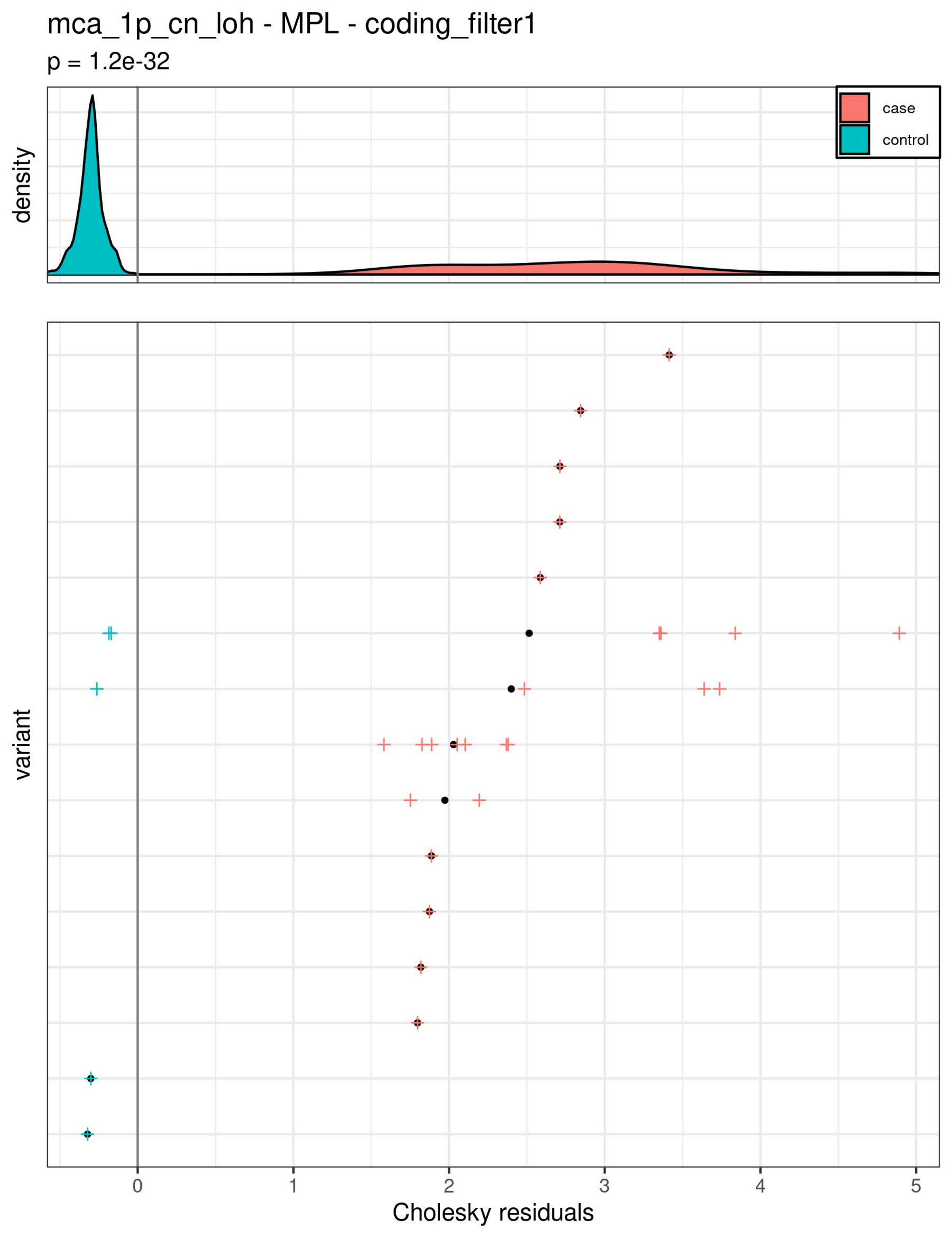

mca_1p_cn_loh - MPL - coding_filter1
p = 1.2e-32
density
case
control
variant
0
1
2
3
4
5
Cholesky residuals

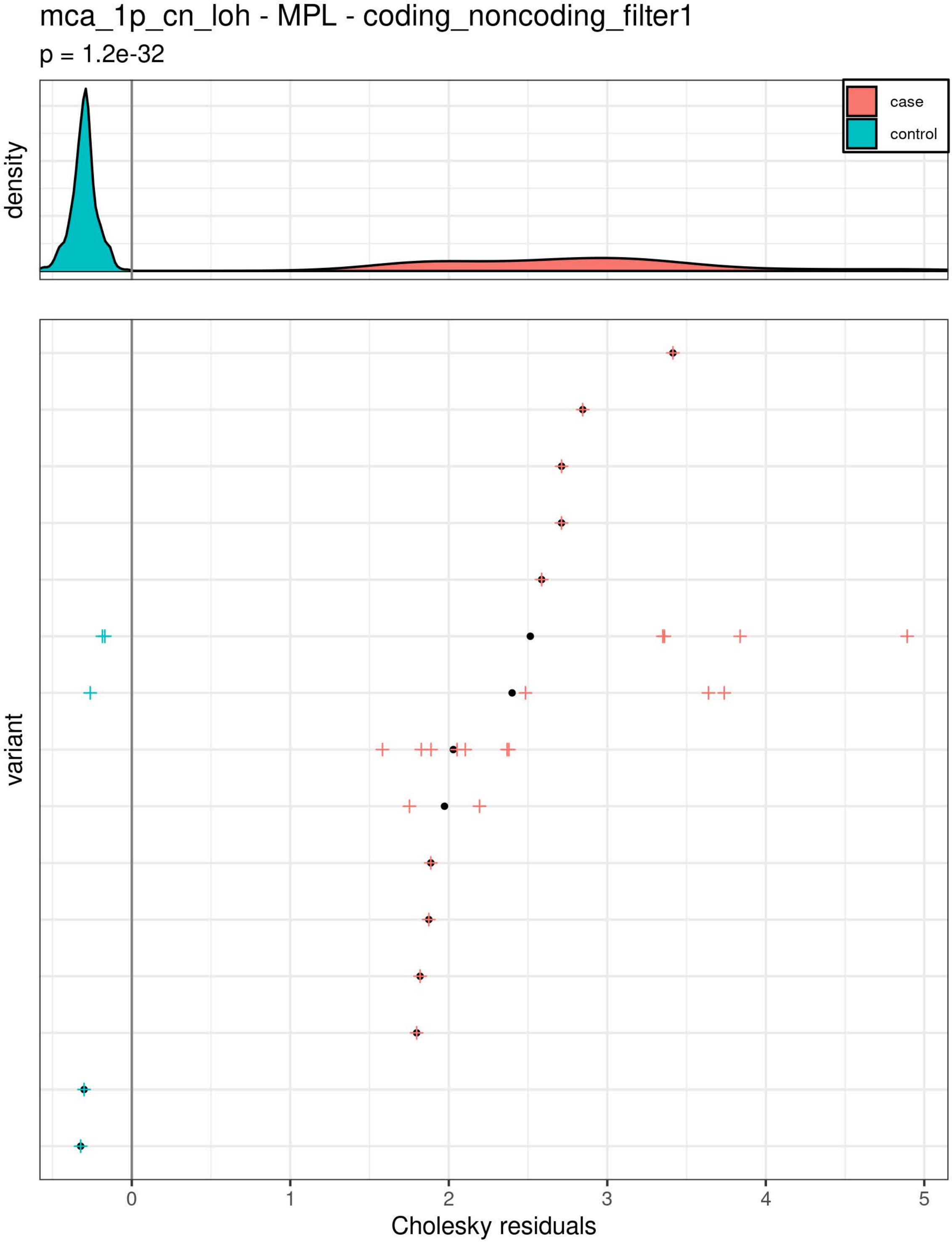
mca_1p_cn_loh - MPL - coding_noncoding_filter1
p = 1.2e-32
density
case
control
variant
0
1
2
3
4
5
Cholesky residuals

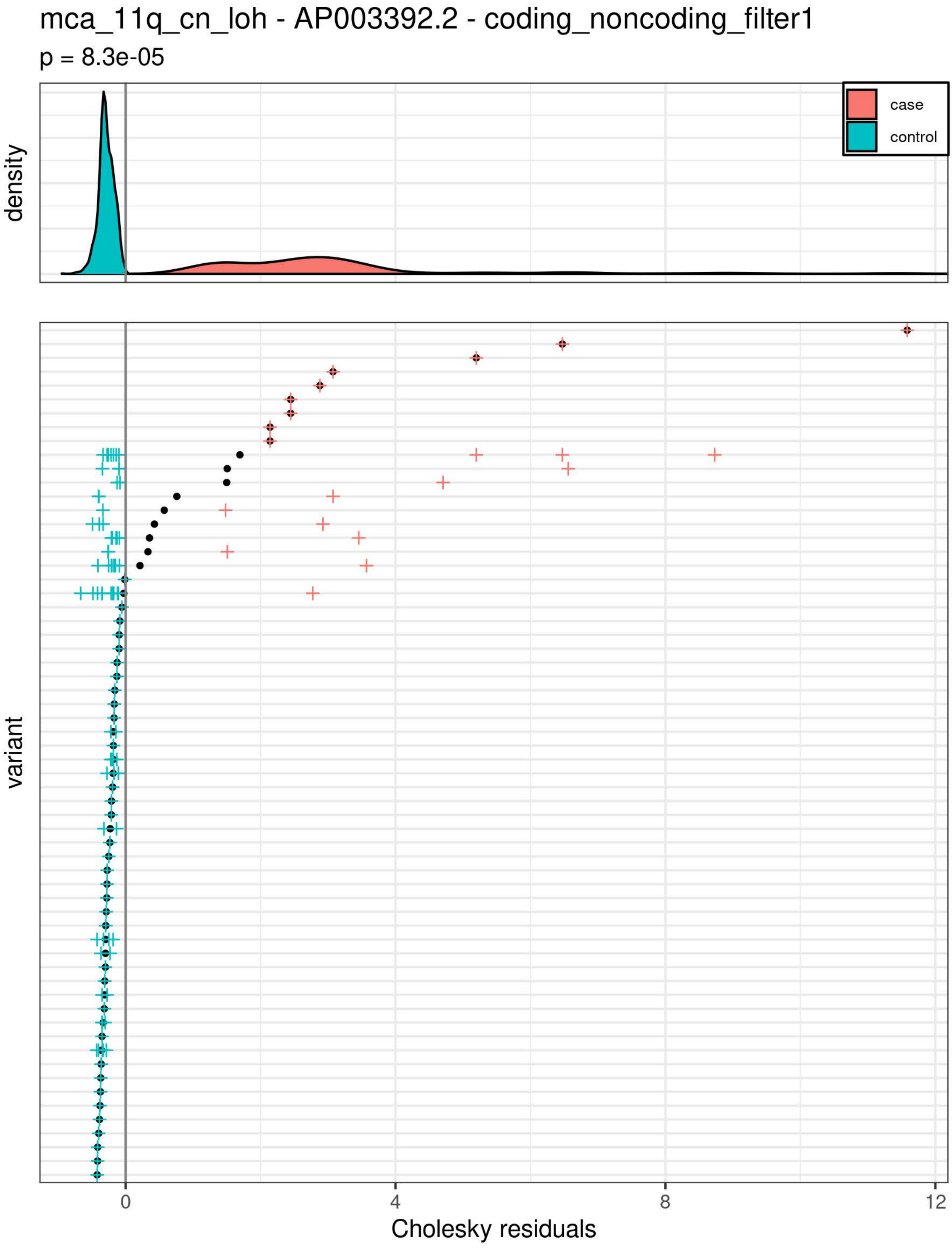

mca_11q_cn_loh - AP003392.2 - coding_noncoding_filter1
p = 8.3e-05
density
case
control
variant
0
4
8
12
Cholesky residuals

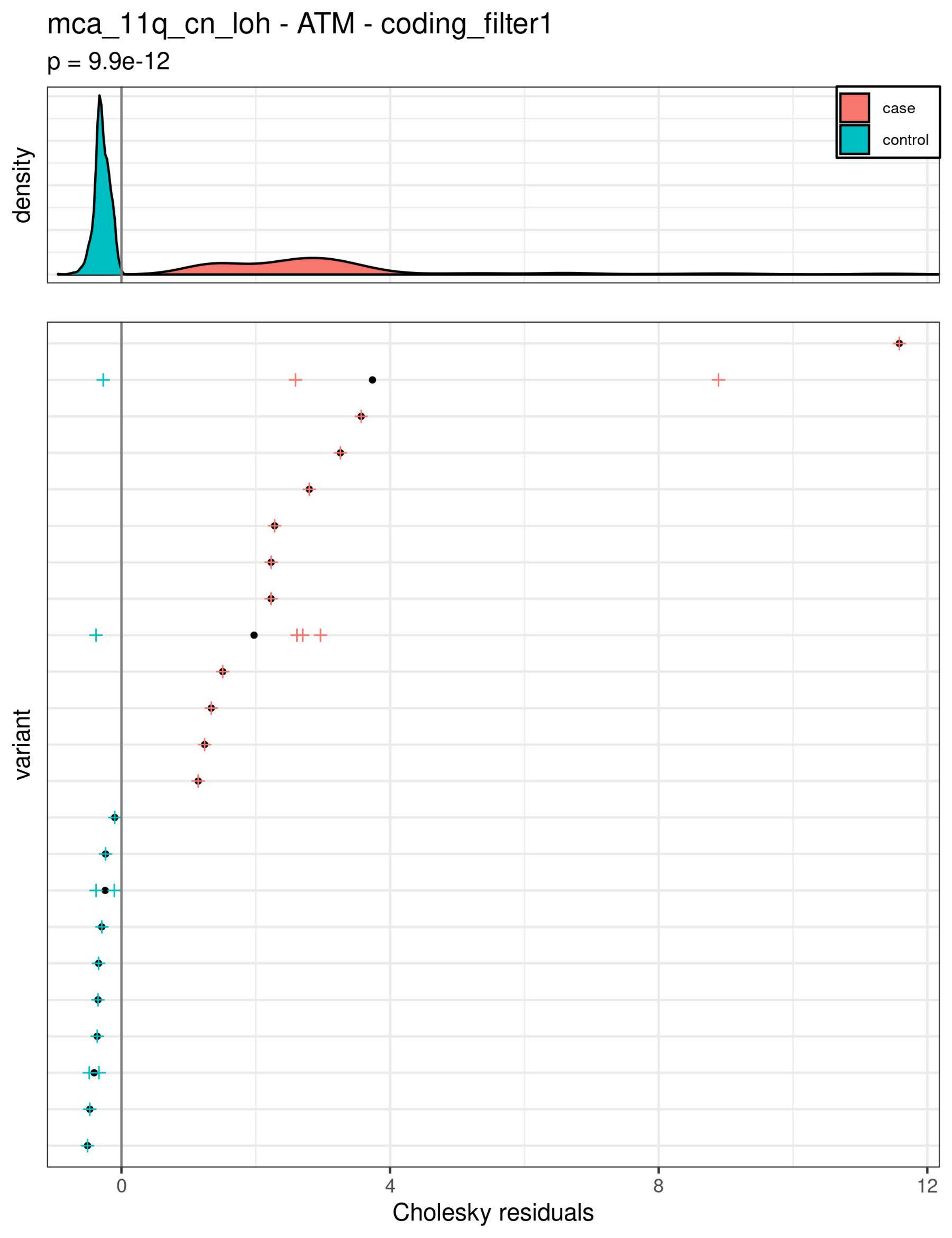

mca_11q_cn_loh - ATM - coding_filter1
p = 9.9e-12
density
case
control
variant
0
4
8
12
Cholesky residuals

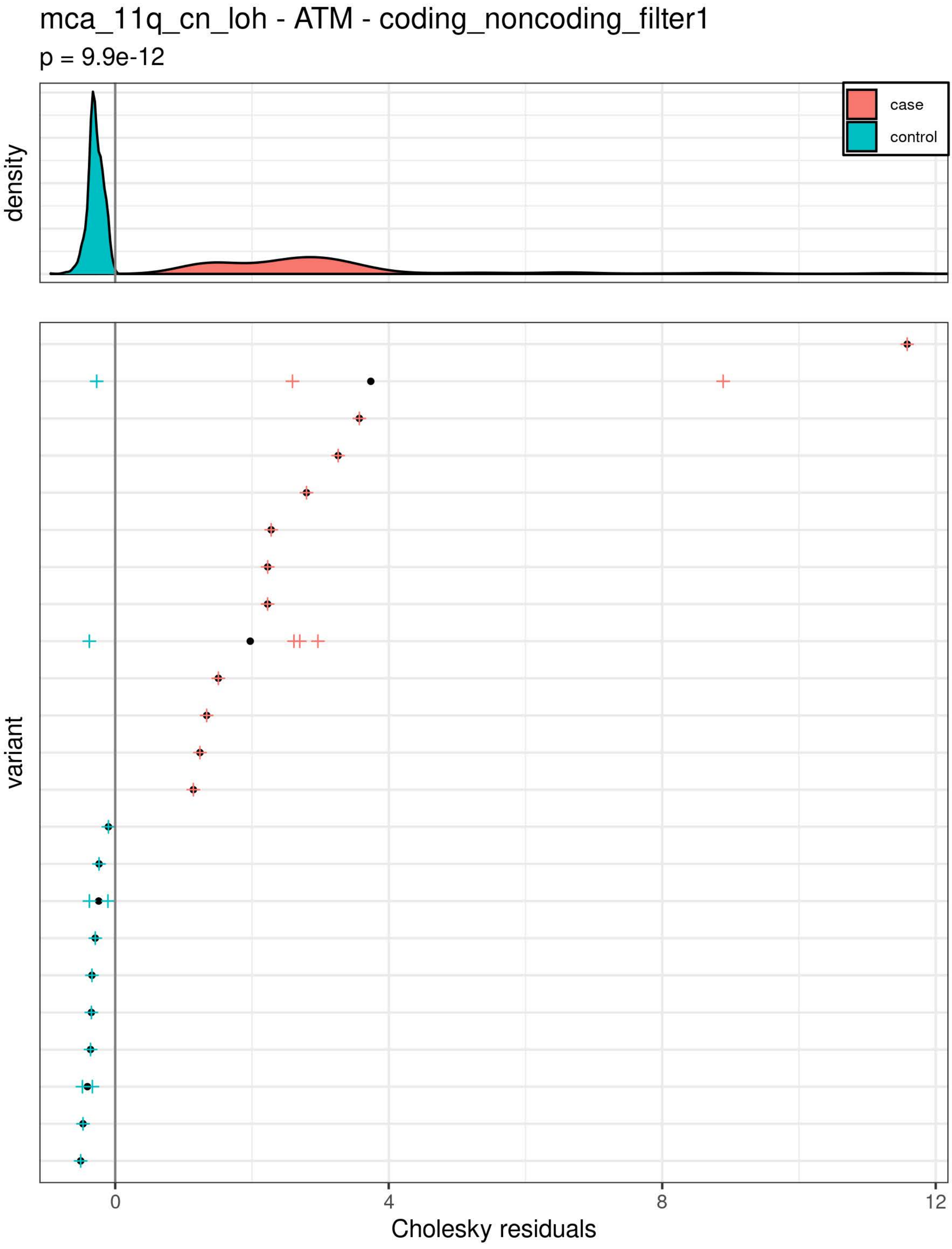

mca_11q_cn_loh - ATM - coding_noncoding_filter1
p = 9.9e-12
density
case
control
variant
0
4
8
12
Cholesky residuals

mca_atm - ATM - coding_noncoding_filter1

p = 1.3e-10

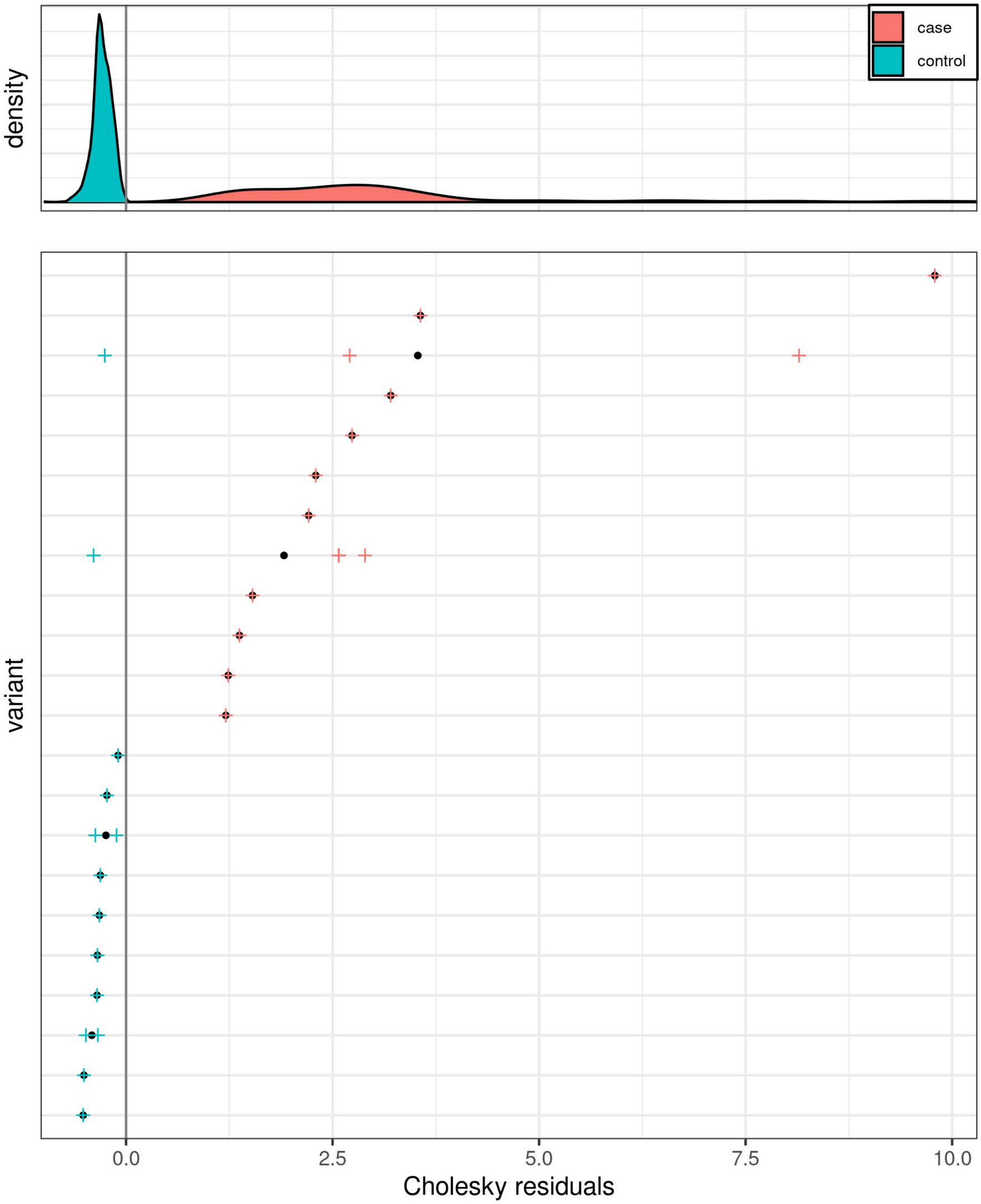

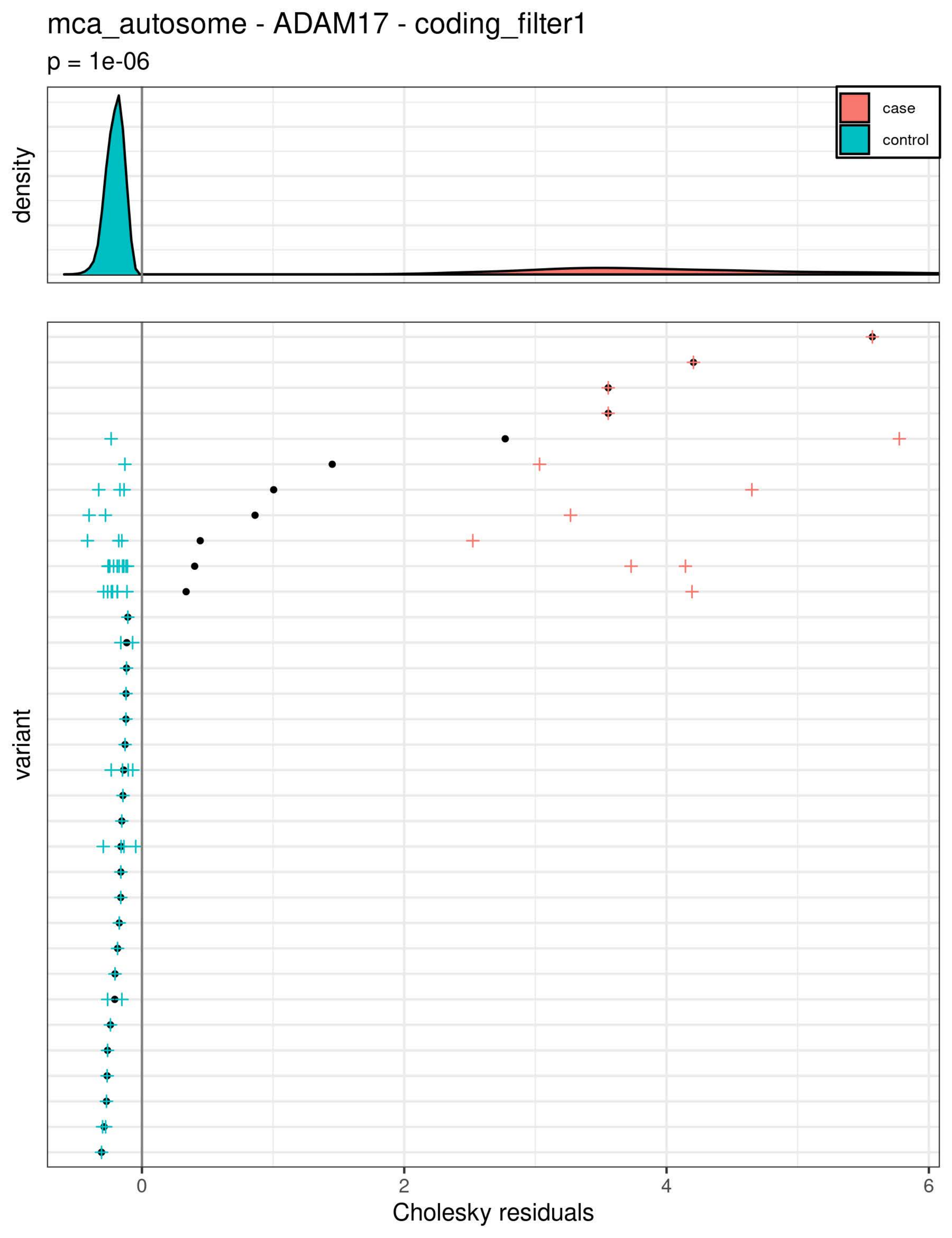

mca_autosome - ADAM17 - coding_filter1
p = 1e-06
density
case
control
variant
0
2
4
6
Cholesky residuals

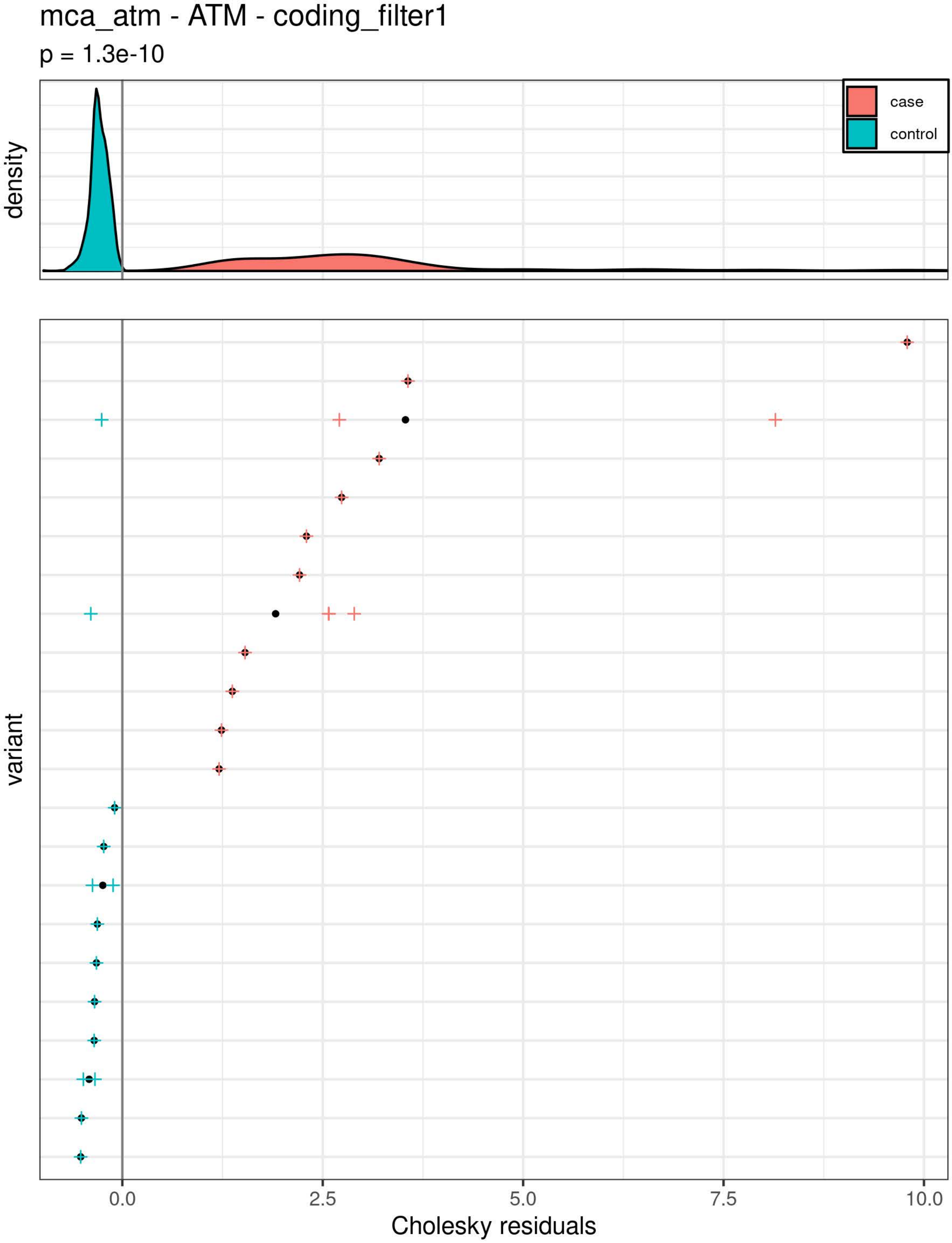
mca_atm - ATM - coding_filter1
p = 1.3e-10
density
case
control
variant
0.0
2.5
5.0
7.5
10.0
Cholesky residuals

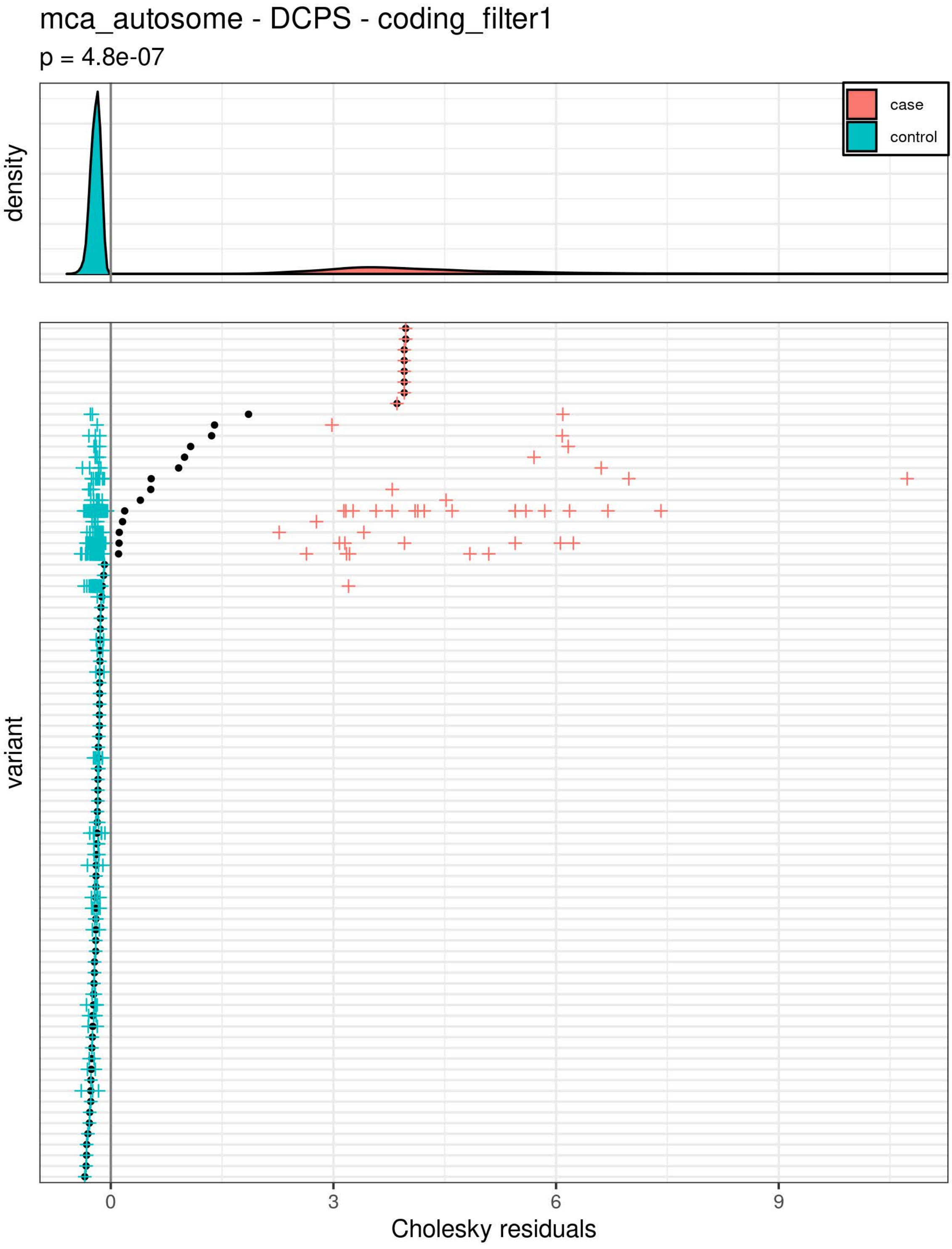
mca_autosome - DCPS - coding_filter1
p = 4.8e-07
density
case
control
variant
0
3
6
9
Cholesky residuals

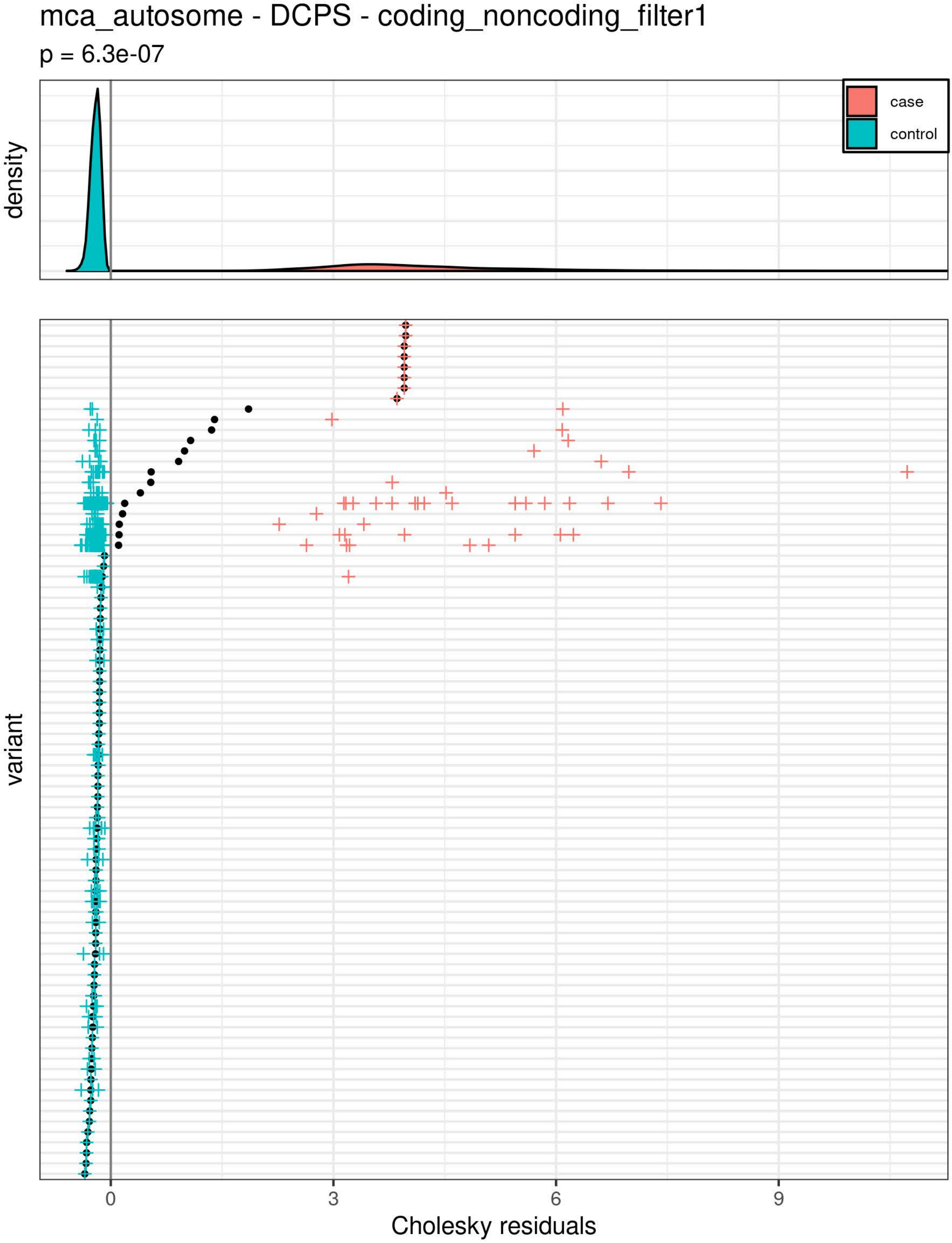
mca_autosome - DCPS - coding_noncoding_filter1
p = 6.3e-07
density
case
control
variant
0
3
6
9
Cholesky residuals

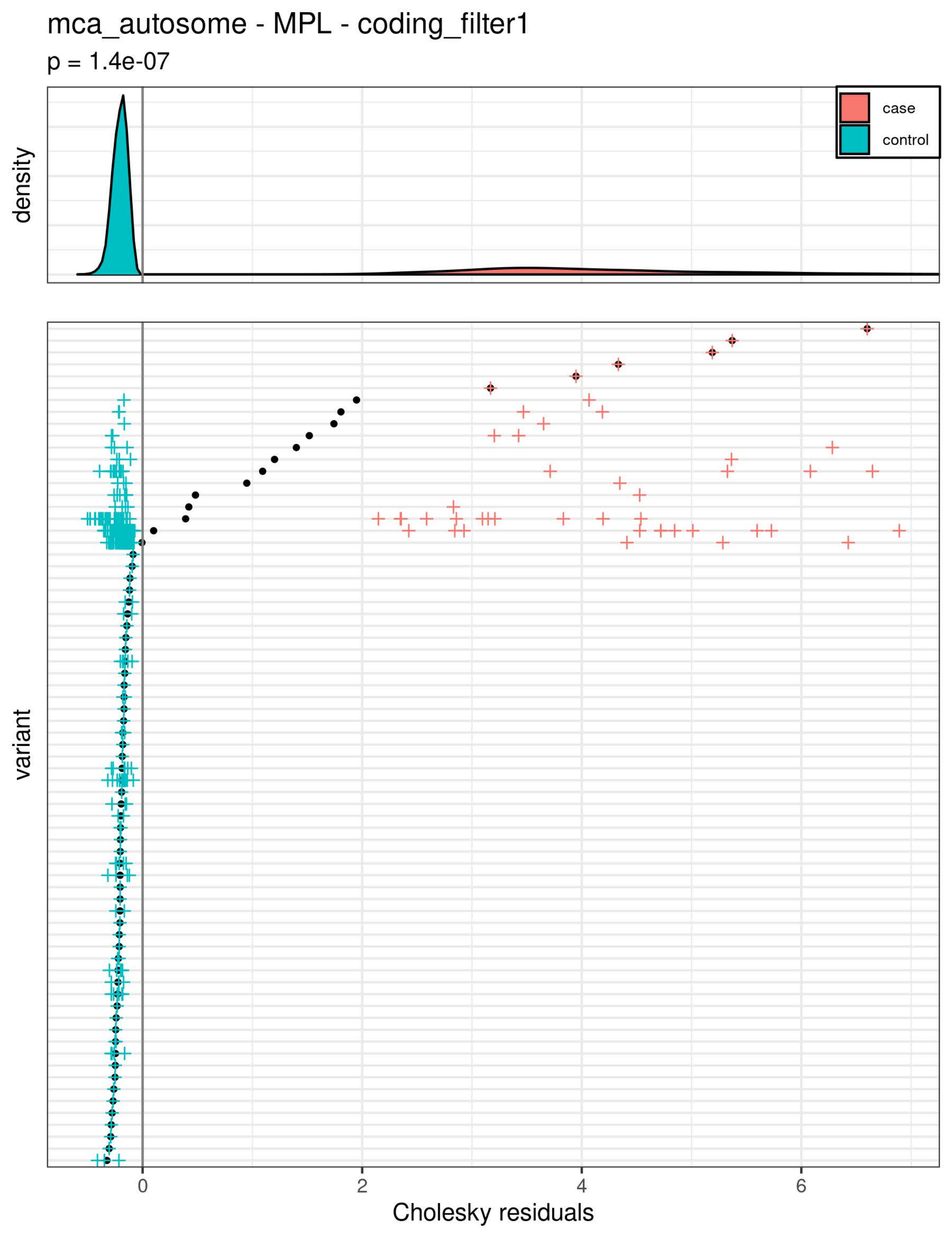
mca_autosome - MPL - coding_filter1
p = 1.4e-07
density
case
control
variant
0
2
4
6
Cholesky residuals

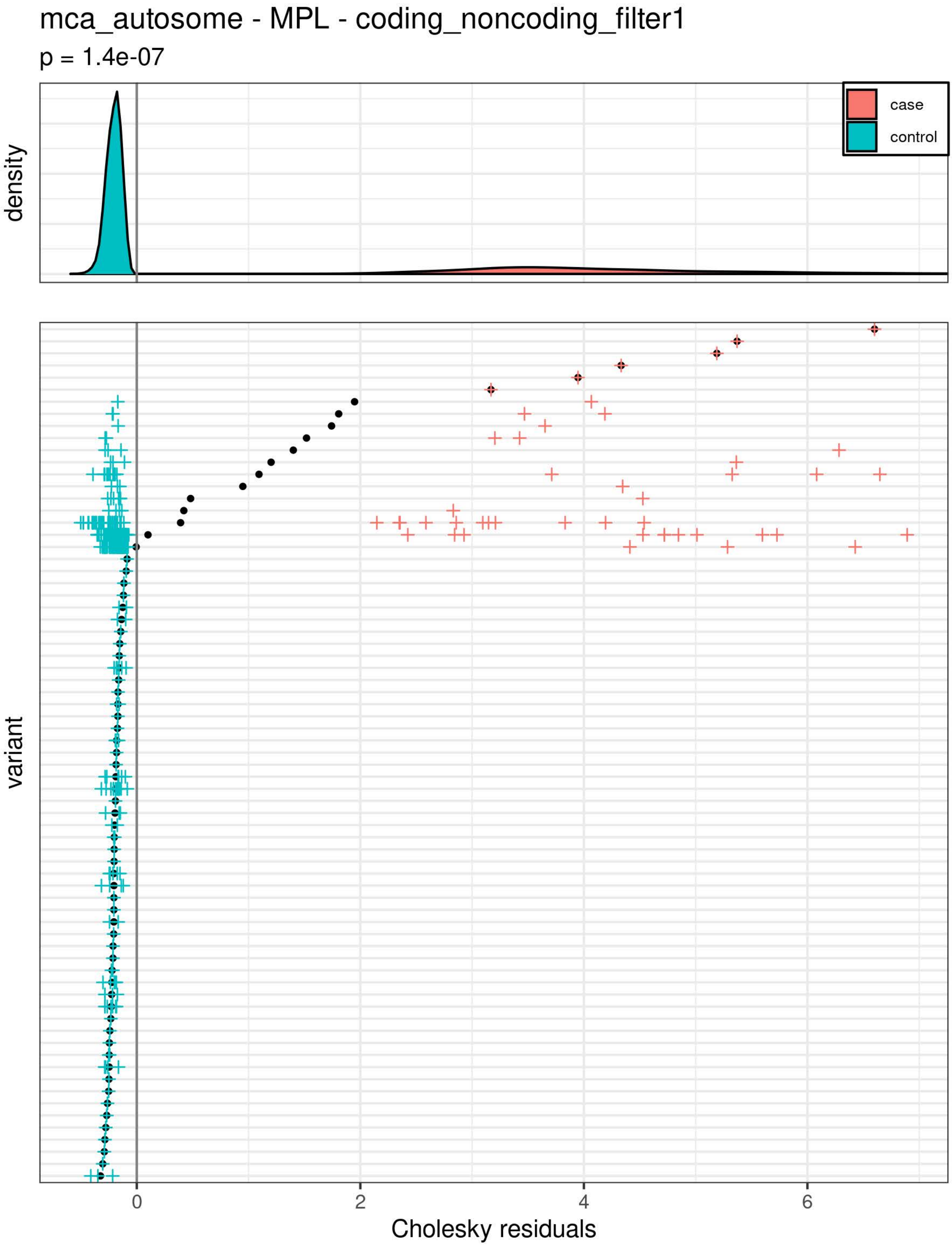
mca_autosome - MPL - coding_noncoding_filter1
p = 1.4e-07
density
case
control
variant
0
2
4
6
Cholesky residuals

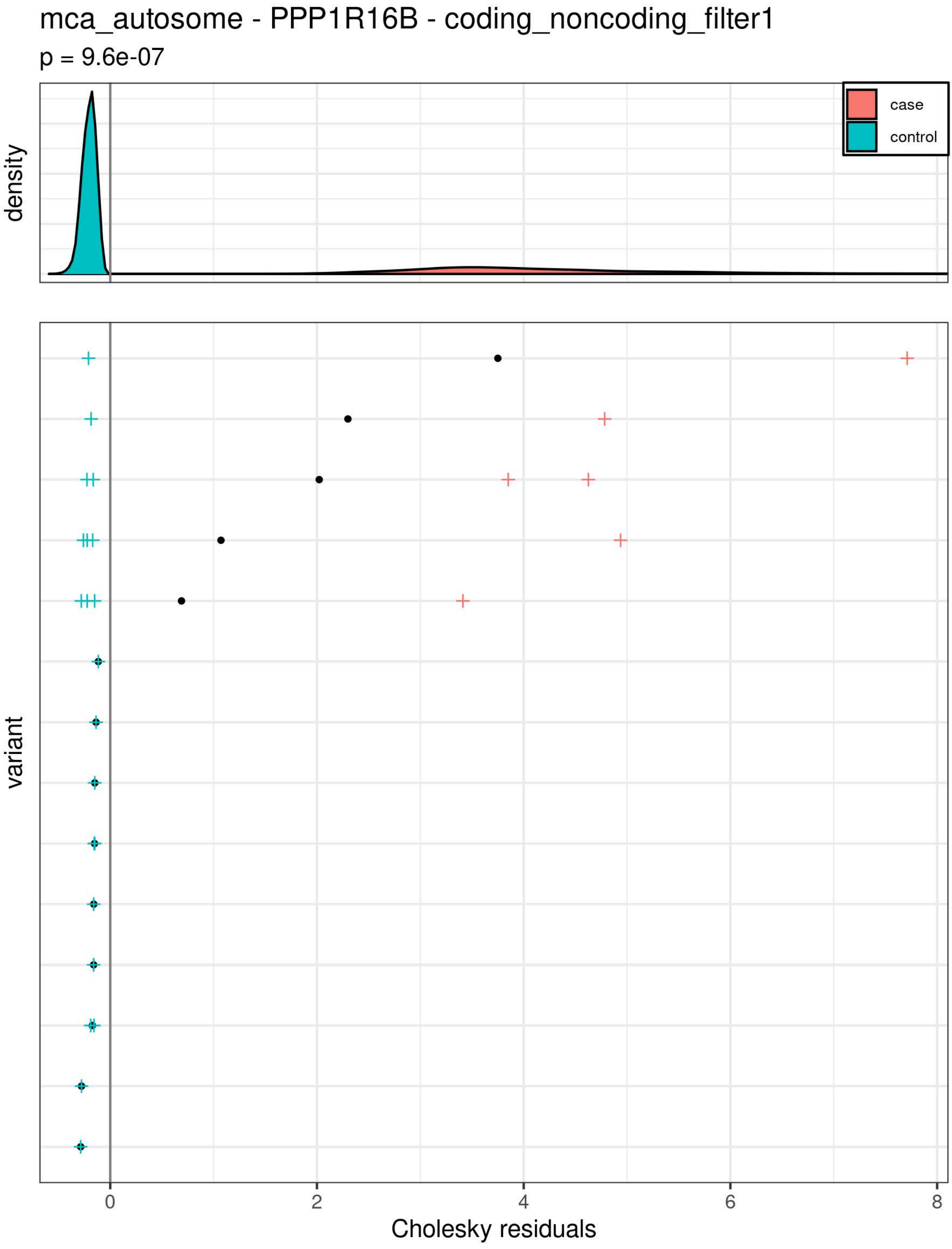
mca_autosome - PPP1R16B - coding_noncoding_filter1
p = 9.6e-07
density
case
control
variant
0
2
4
6
8
Cholesky residuals

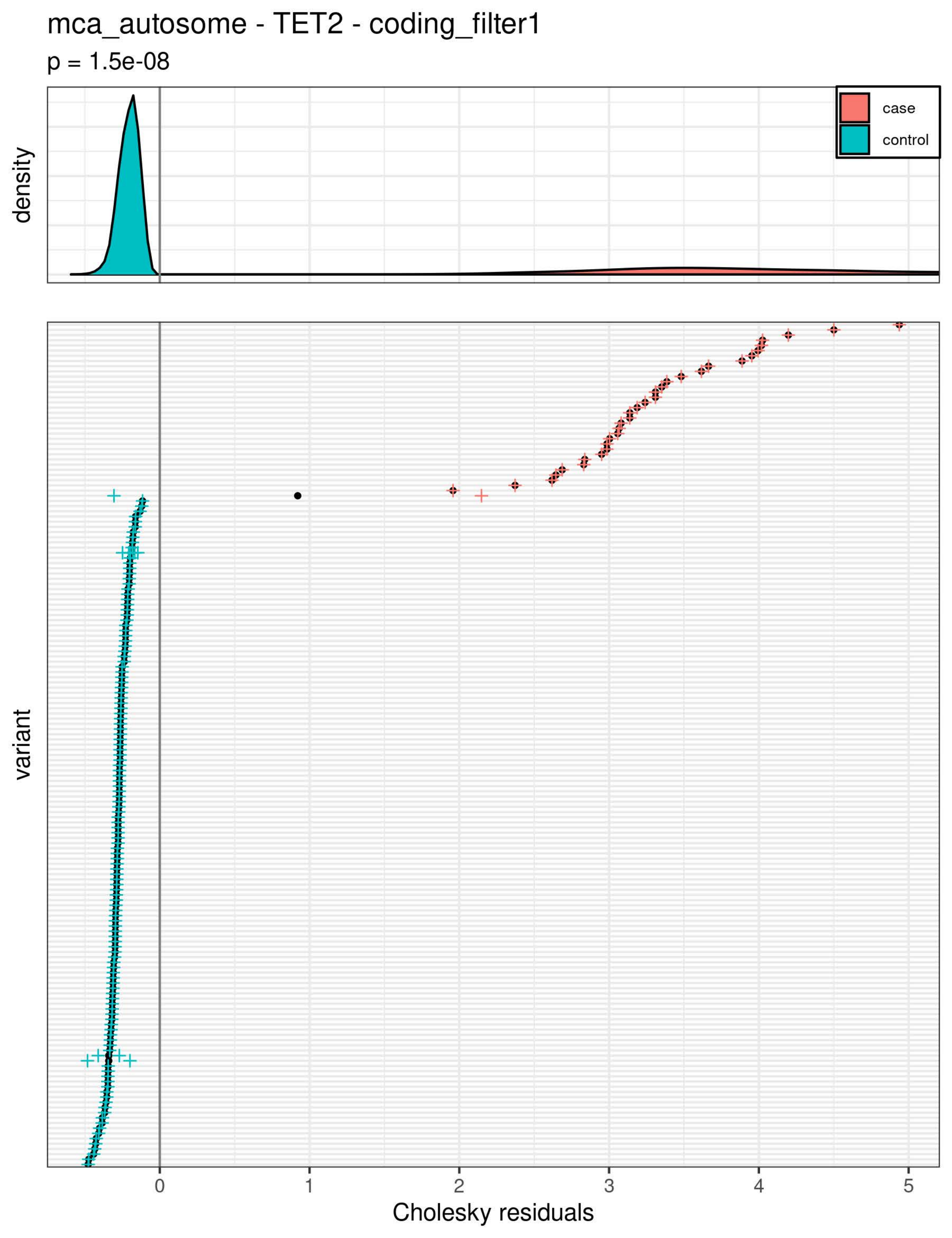
mca_autosome - TET2 - coding_filter1
p = 1.5e-08
density
case
control
variant
0
1
2
3
4
5
Cholesky residuals

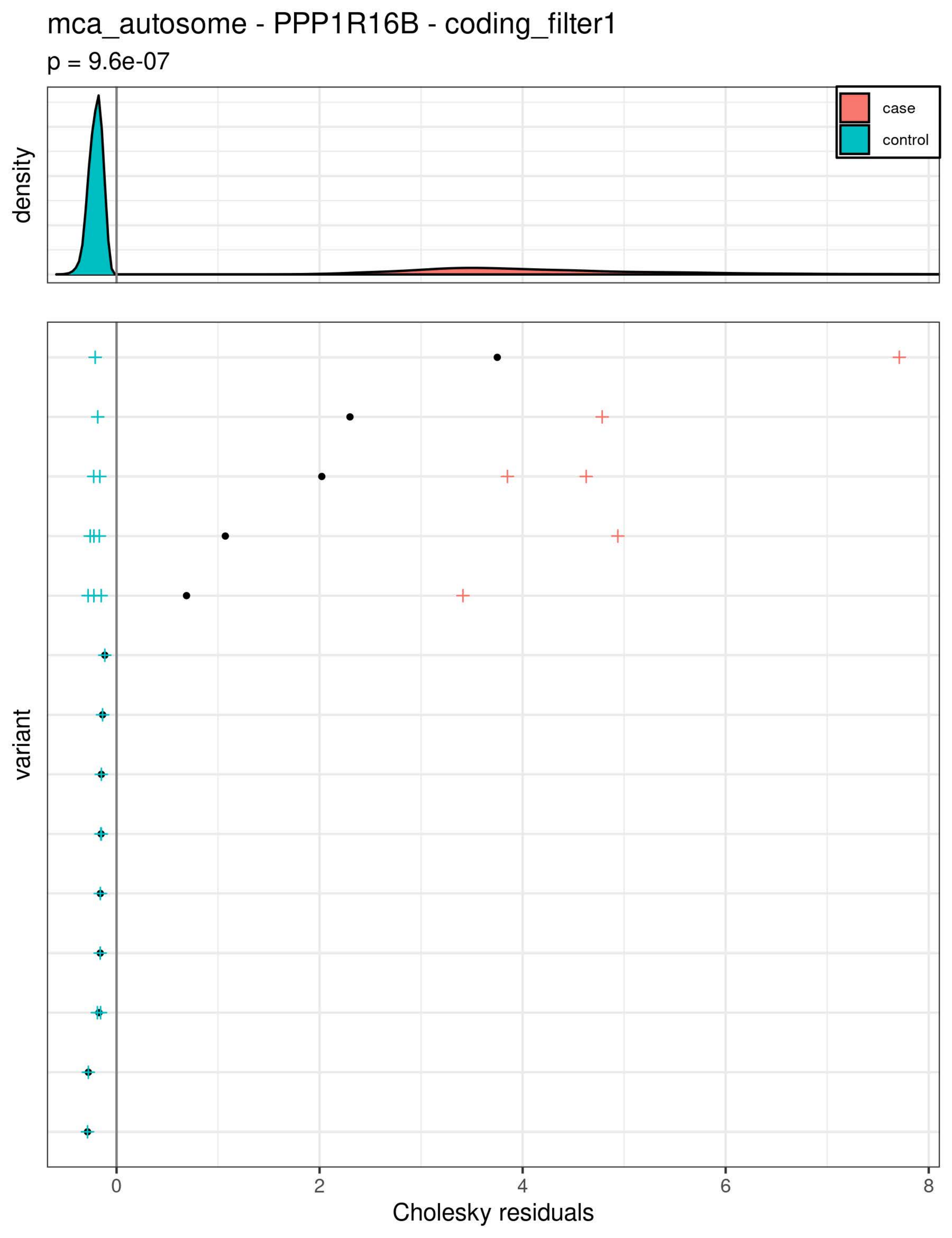
mca_autosome - PPP1R16B - coding_filter1
p = 9.6e-07
density
case
control
variant
0
2
4
6
8
Cholesky residuals

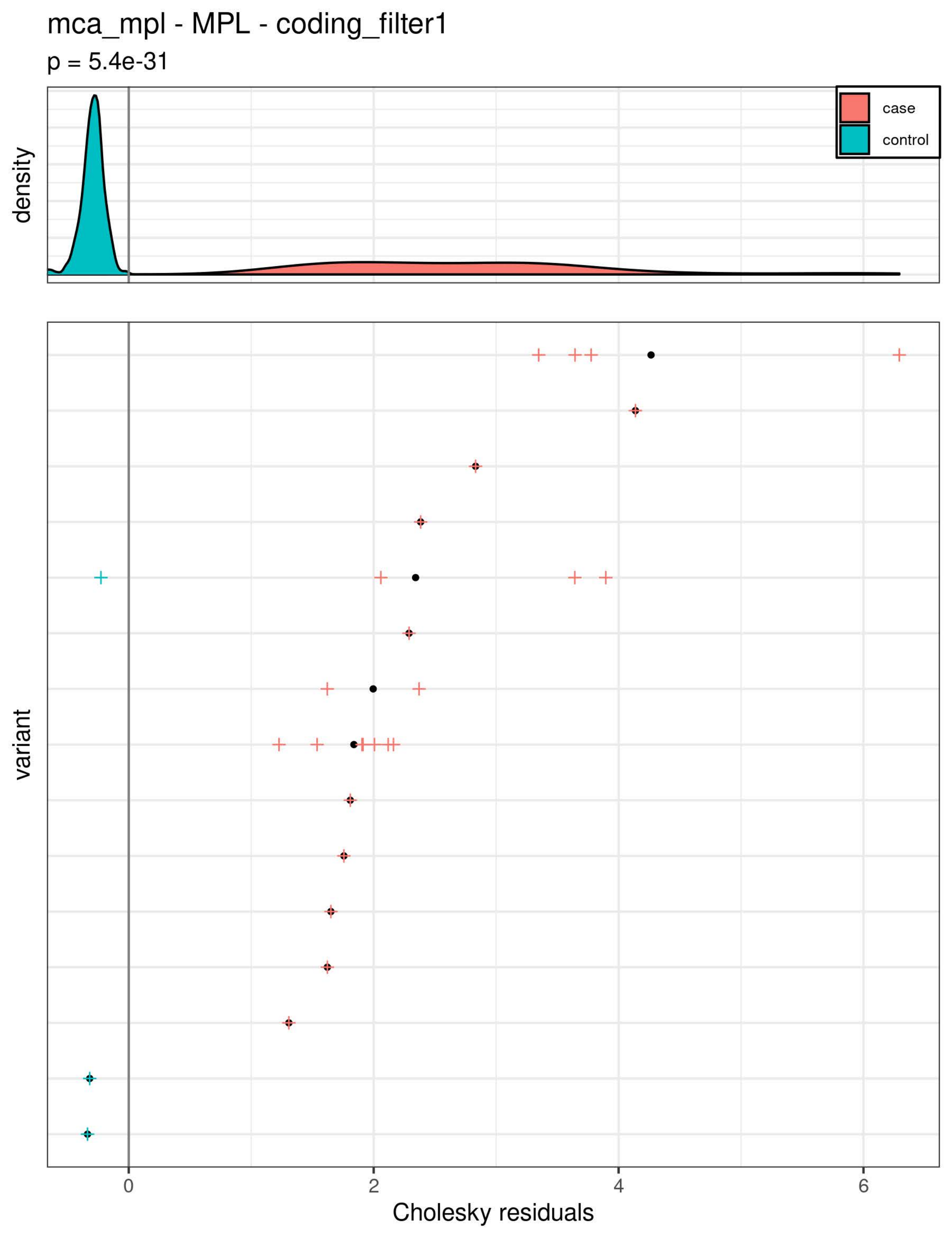
mca_mpl - MPL - coding_filter1
p = 5.4e-31
density
case
control
variant
0
2
4
6
Cholesky residuals

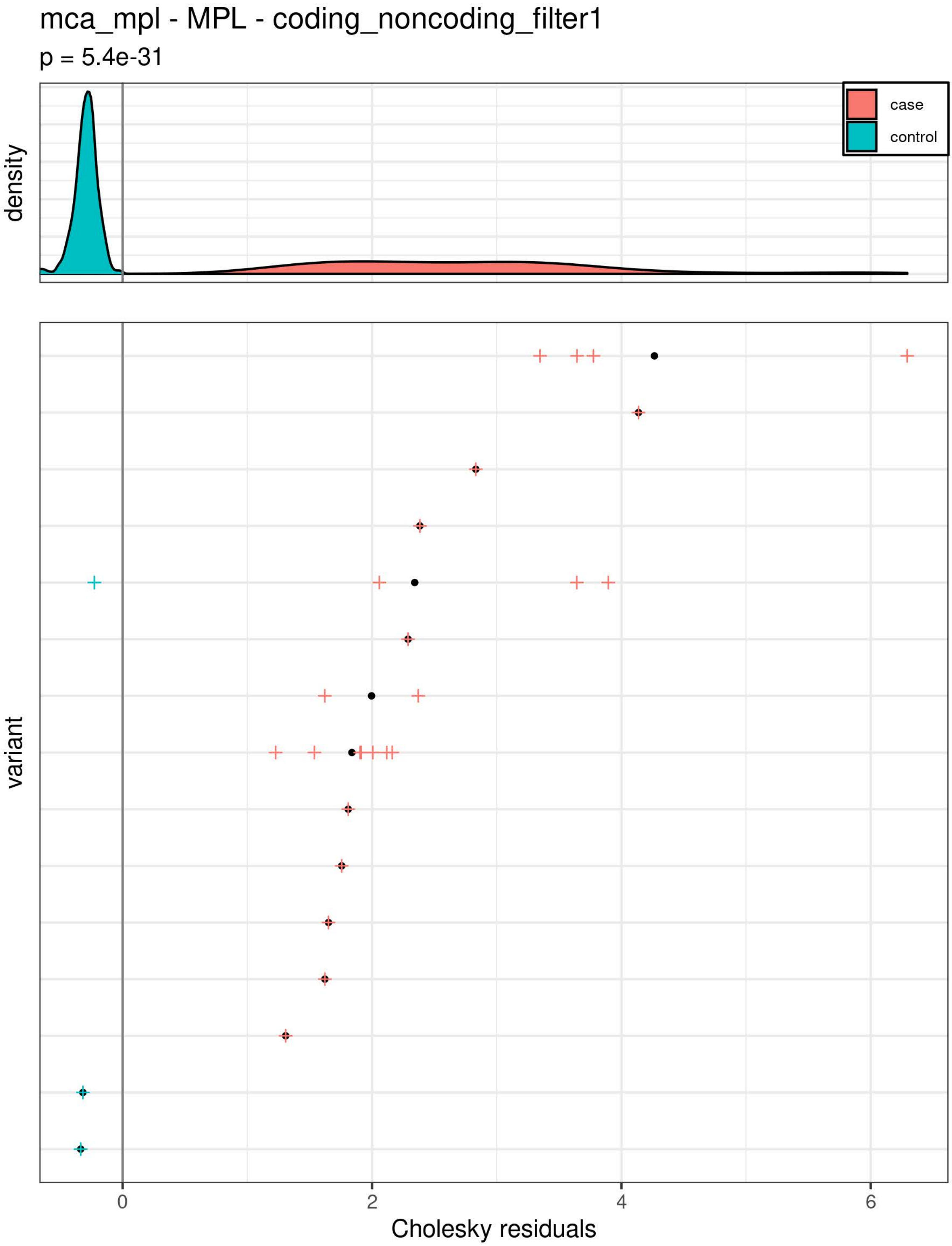
mca_mpl - MPL - coding_noncoding_filter1
p = 5.4e-31
density
case
control
variant
0
2
4
6
Cholesky residuals

mca_x - OR4C16 - coding_filter1

p = 2.4e-06

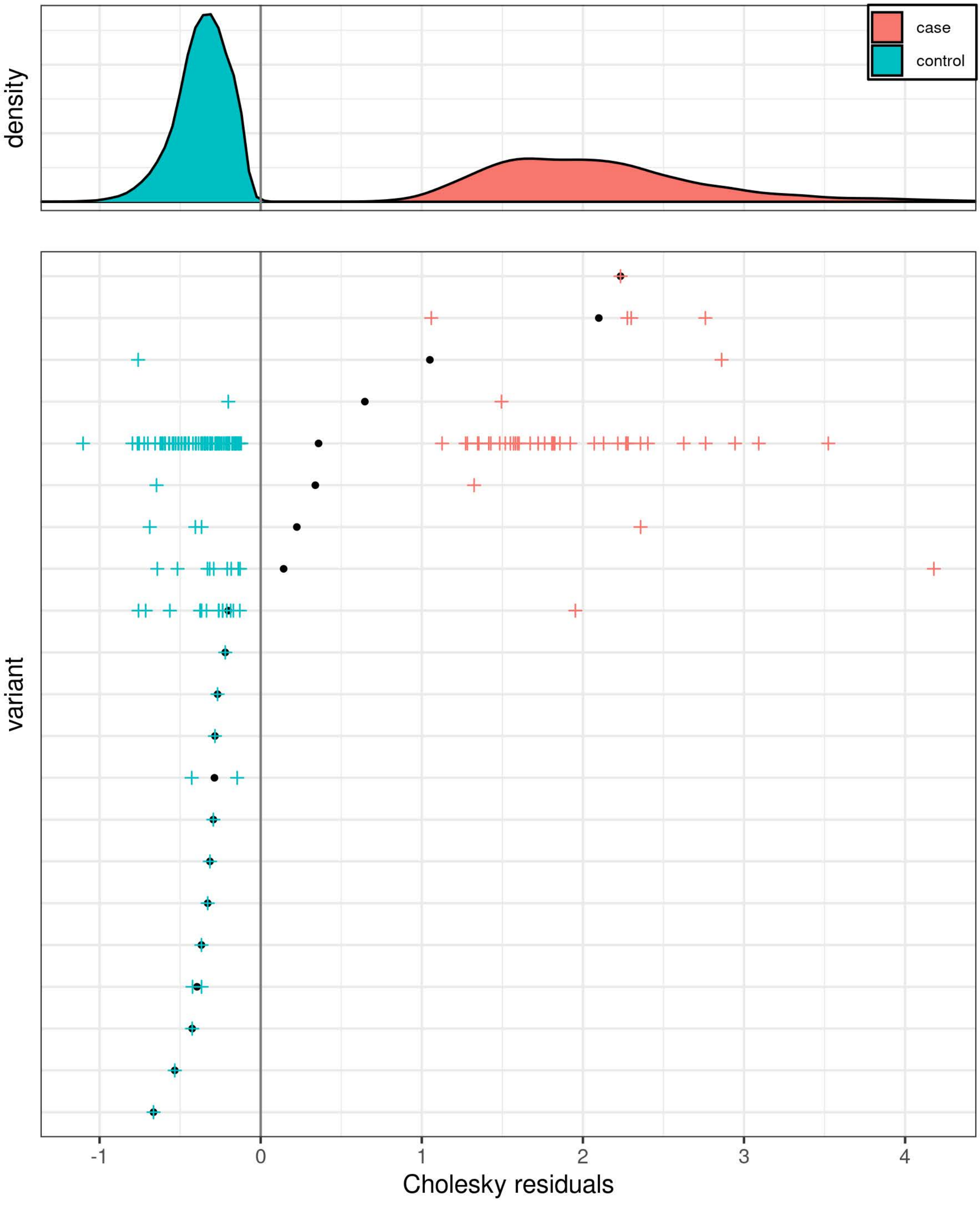
