## Supplemental Methods for "Mosaic chromosomal alterations in blood across ancestries via whole-genome sequencing"

**Supplementary Materials**

**Participating TOPMed studies**

*Amish*

The Amish Complex Disease Research Program includes a set of large community-based studies focused largely on cardiometabolic health carried out in the Old Order Amish (OOA) community of Lancaster, Pennsylvania (http://medschool.umaryland.edu/endocrinology/amish/research-program.asp). The OOA population of Lancaster County, PA immigrated to the Colonies from Western Europe in the early 1700’s. There are now over 30,000 OOA individuals in the Lancaster area, nearly all of whom can trace their ancestry back 12-14 generations to approximately 700 founders. Investigators at the University of Maryland School of Medicine have been studying the genetic determinants of cardiometabolic health in this population since 1993. To date, over 7,000 Amish adults have participated in one or more of our studies.

Due to their ancestral history, the OOA are enriched for rare exonic variants that arose in the population from a single founder (or small number of founders) and propagated through genetic drift. Many of these variants have large effect sizes and identifying them can lead to new biological insights about health and disease. The parent study for this WGS project provides one (of multiple) examples. In our parent study, we identified through a genome-wide association analysis a haplotype that was highly enriched in the OOA that is associated with very high LDL-cholesterol levels. At the present time, the identity of the causative SNP – and even the implicated gene – is not known because the associated haplotype contains numerous genes, none of which are obvious lipid candidate genes. A major goal of the WGS that will be obtained through the NHLBI TOPMed Consortium will be to identify functional variants that underlie some of the large effect associations observed in this unique population.

*ARIC*

The ARIC study is a population-based prospective cohort study of cardiovascular disease sponsored by the National Heart, Lung, and Blood Institute (NHLBI). ARIC included 15,792 individuals, predominantly European American and African American, aged 45-64 years at baseline (1987-89), chosen by probability sampling from four US communities. Cohort members completed three additional triennial follow-up examinations, a fifth exam in 2011-2013, a sixth exam in 2016-2017, and a seventh exam in 2018-2019. The ARIC study has been described in detail previously (The ARIC Investigators. The Atherosclerosis Risk in Communities (ARIC) study: Design and objectives. American Journal of Epidemiology 1989;129:687-702).

*BAGS*

Epidemiologic studies of asthma have been underway in Barbados since 1991, when PI Barnes reported a relationship between modernization of the domestic environment in Barbados and increased risk of asthma. The baseline prevalence of asthma in Barbados is high (~20%), and from admixture analyses, we have determined that the proportion of African ancestry among Barbadian founders is similar to U.S. African Americans, rendering this a unique population to disentangle the genetic basis for asthma disparities among African ancestry populations in general. The primary outcome measure is asthma, and the approach for characterizing asthma in the Barbados population is based on the validated Respiratory Health Questionnaire (RHQ) designed from the 1978 American Thoracic Society questionnaire. Additional phenotype data include lung function measures, asthma severity, total serum IgE, and serum levels of various cytokines. In 1993, the *Barbados Asthma Genetics Study* (BAGS) was initiated on nuclear and extended asthmatic families who self-reported as African Caribbean, resulting in the first evidence for linkage for asthma and tIgE in an African-ancestry population, and the development of novel family-based methods. Recruitment into the BAGS program was enhanced through its involvement in the international *Genetics of Asthma International Network* (1999-2001) and the current sample of >1300 participants continues to grow through the efforts of collaborators and nursing staff at the Chronic Disease Research Centre in Barbados. Pediatric probands were recruited through referrals at local polyclinics or the Accident and Emergency Department at the Queen Elizabeth Hospital, and their nuclear and extended family members were subsequently recruited. All subjects gave verbal and written consent as approved by the Johns Hopkins Institutional Review Board (IRB) and the Barbados Ministry of Health.

In 2007 we performed a genome-wide association study (GWAS) on 655,352 SNPs using the Illumina Infinium™ II HumanHap650Y BeadChip v.1.0 (Illumina Inc.) on a subset of 1,000 Barbados participants. This represented the first GWAS of asthma focusing exclusively on populations of African ancestry, and data from this study also contributed to the NHLBI-supported EVE Consortium. BAGS also contributed 96 samples to Phase 2 of the Thousand Genomes Project (TGP). Subsequently, BAGS samples were included in the NHLBI-supported parent grant, entitled *New Approaches for Empowering Studies of Asthma in Populations of African Descent*“ (R01 HL104608-01), in which whole genome sequencing (WGS) was performed on ~1,000 individuals from North, Central, and South American and Caribbean and two West African populations. These populations constitute the *Consortium on Asthma among African-ancestry Populations in the Americas* (CAAPA), which aims to discover genes influencing risk for asthma, and catalog genetic diversity in descendants of the African Diaspora in the Americas. So far, CAAPA sequencing has greatly expanded the lexicon of human diversity, as we have observed >20% more variants than reported in the 1000 Genome Project (TGP). Using these WGS data, a custom, gene-centric SNP genotyping array was developed by Illumina, Inc., called the *African Diaspora Power Chip* (ADPC), to complement current, commercially available genome-wide chips, which provide sub-optimal tagging of genes among individuals of African ancestry. This ADPC was recently genotyped on all BAGS samples, with a goal of combining ADPC data with existing GWAS data from the 650Y to test for association with asthma. The initial goals of the parent grant did not include validating the ADPC. Moreover, the ADPC, combined with existing GWAS data, will be limited in detecting contributions of rare and structural variants, which may account for some of the “missing heritability” of asthma. We therefore are performing WGS on 1,100 asthmatics and family members from the BAGS, in order to (i) expand the CAAPA WGS dataset and thereby the genomic catalog of African ancestry for the research community; (ii) validate the ADPC by capturing information from both common and rare variants; and (iii) generate additional discovery of rare and structural variants that may control risk to asthma. Tools resulting from this study will result in substantial advancements in the technology available for identifying genes relevant to disease in under-represented minorities.

Given the data available on this large, deeply genotyped cohort from a relatively homogeneous environment representing an underrepresented minority group suffering most from asthma, the BAGS sample provides a unique opportunity to employ novel genomics.

*BioMe*

The Charles Bronfman Institute for Personalized Medicine at Mount Sinai Medical Center (MSMC), BioMe Biobank, founded in September 2007, is an ongoing, broadly-consented electronic health record-linked clinical care biobank that enrolls participants non-selectively from the Mount Sinai Medical Center patient population. The MSMC serves diverse local communities of upper Manhattan, including Central Harlem (86% African American), East Harlem (88% Hispanic/Latino), and Upper East Side (88% Caucasian/White) with broad health disparities.

*CARDIA*

The Coronary Artery Risk Development in Young Adults (CARDIA) Study is a study examining the development and determinants of clinical and subclinical cardiovascular disease and their risk factors. It began in 1985-1986 with a group of 5,115 black and white men and women aged 18-30 years. The participants were selected so that there would be approximately the same number of people in subgroups of race, gender, education (high school or less and more than high school) and age (18-24 and 25-30) in each of 4 centers: Birmingham, AL; Chicago, IL; Minneapolis, MN; and Oakland, CA.

*CFS*

The CFS is a genetic epidemiological study of 352 rigorously phenotyped families ascertained through probands with OSA identified through Cleveland, OH area sleep centers, neighborhood controls, and the spouses and first and second degree relatives of probands. Participants were studied on up to 4 exams between 1990-2006 with overnight sleep studies, standardized anthropometry; questionnaires; blood pressure; and spirometry. Fasting serum and ECGs are available from the last exam. Participants have a mean age of 37.7 years (African Americans) and 41.4 years (European Americans). Slightly more than 50% of the sample is female and 31% have moderate to severe OSA; 12.6% have diabetes, and 34.0% have hypertension. Asthma is reported in 19% and 13% of the African Americans and European Americans, respectively. Heritability analysis of traditional OSA traits as well as novel traits such as hypopnea duration (a marker of respiratory arousability) as well as overnight oxygenation (a marker of susceptibility to hypoxemia occurring with recurrent apneas) has shown that the latter traits are more heritable (h2 > 0.50) than traditional measures. Linkage analysis has identified peaks (and individual families contributing to peaks) for these traits. Through the Life After Linkage initiative (5R01HL113338), we further have aggregated and analyzed data on 19,798 individuals from 7 cohorts (Cleveland Family Study [CFS] plus ARIC, FHS, HCHS/SOL, MESA, MrOS, and Starr County) and conducted the largest GWAS to date of OSA traits.

*CHS*

The Cardiovascular Health Study (CHS) is a population-based cohort study of risk factors for coronary heart disease and stroke in adults 65 years and older conducted across four field centers 2. The original predominantly European ancestry cohort of 5,201 persons was recruited in 1989-1990 from random samples of people on Medicare eligibility lists from four US communities. Subsequently, an additional predominantly African-American cohort of 687 persons was enrolled for a total sample of 5,888. Institutional review committees at each field center approved the CHS, and participants gave informed consent. Blood samples were drawn from all participants at their baseline examination, and DNA was subsequently extracted from available samples. These analyses were limited to participants with available DNA who also consented to genetic studies. Participants were examined annually from enrollment to 1999 and continued to be under surveillance for stroke following 1999.

*COPDGene*

COPDGene (also known as the Genetic Epidemiology of COPD Study) is an NIH-funded, multicenter study. A study population of more than 10,000 smokers (1/3 African American and 2/3 non-Hispanic White) has been characterized with a study protocol including pulmonary function tests, chest CT scans, six minute walk testing, and multiple questionnaires. Five and ten years after this initial visit, all available study participants are being brought back for a follow-up visit with a similar study protocol. This study has been used for epidemiologic and genetic studies. Previous genetic analysis in this study has been based on genome-wide SNP genotyping data. More than 10,000 subjects underwent whole genome sequencing in this NHLBI WGS project, including severe COPD subjects and resistant smoking controls. The COPDGene Study web site is: http://www.copdgene.org/.

*FHS*

FHS is a three-generation, single-site, community-based, ongoing cohort study that was initiated in 1948 to investigate prospectively the risk factors for CVD including stroke. It now comprises 3 generations of participants: the Original cohort followed since 1948; their Offspring and spouses of the Offspring, followed since 1971 4; and children from the largest Offspring families enrolled in 2002 (Gen 3). The Original cohort enrolled 5,209 men and women who comprised two-thirds of the adult population then residing in Framingham, MA. Survivors continue to receive biennial examinations. The Offspring cohort comprises 5,124 persons (including 3,514 biological offspring) who have been examined approximately once every 4 years. The Gen 3 cohort contains 4,095 participants.

*GeneSTAR*

In 1982 The Johns Hopkins Sibling and Family Heart Study was created to study patterns of coronary heart disease and related risk factors in families with early-onset coronary disease, identified from10 Baltimore area hospitals. Renamed in 2003, the Genetic Study of Atherosclerosis Risk (GeneSTAR) continues to study mechanisms of coronary heart disease and stroke in families using novel models and exciting new methods. GeneSTAR is a family-based study including initially healthy brothers and sisters identified from probands with early-onset coronary disease, along with the healthy offspring of the siblings and the probands. The goal is to discover and amplify mechanisms of stroke and coronary heart disease. Our African American and European American family cohort has undergone extensive screening, genetic testing, and follow-up for new cardiovascular disease, stroke, and other clinical events for 5 to 38 years.

*HCHS/SOL*

The Hispanic Community Health Study/Study of Latinos (HCHS/SOL) is a multi-center study of Hispanic/Latino populations with the goal of determining the role of acculturation in the prevalence and development of diseases, and to identify other traits that impact Hispanic/Latino health. The study is sponsored by the National Heart, Lung, and Blood Institute (NHLBI) and other institutes, centers, and offices of the National Institutes of Health (NIH). Recruitment began in 2006 with a target population of 16,000 persons of Cuban, Puerto Rican, Dominican, Mexican or Central/South American origin. Participants were recruited through four sites affiliated with San Diego State University, Northwestern University in Chicago, Albert Einstein College of Medicine in Bronx, New York, and the University of Miami. Recruitment was implemented through a two-stage area household probability design. The study enrolled 16,415 participants who were self-identified Hispanic/Latino and aged 18-74 years, and the extensive psycho-social and clinical assessments were conducted during 2008-2011. Annual telephone follow-up interviews are ongoing since study inception. During the 2014-2017 second visit, the participants were re-examined again for various health outcomes of interest.

*GENOA*

The Genetic Epidemiology Network of Arteriopathy (GENOA) is one of four networks in the NHLBI Family-Blood Pressure Program (FBPP). The long-term objective of GENOA is to elucidate the genetics of target organ complications of hypertension, including both

atherosclerotic and arteriosclerotic complications involving the heart, brain, kidneys, and

peripheral arteries. The longitudinal GENOA study enrolled sibships in which at least two siblings had clinically diagnosed essential hypertension before age 60 years. All other members of the sibship were invited to participate regardless of their hypertension status. Participants were diagnosed with hypertension if they had either 1) a previous clinical diagnosis of hypertension by a physician with current anti-hypertensive treatment, or 2) an average systolic blood pressure ≥ 140 mm Hg or diastolic blood pressure ≥ 90 mm Hg based on the second and third readings at the time of their clinic visit. The initial exam (1995-2000) enrolled 1583 non-Hispanic white participants from Rochester, Minnesota, and 1854 African American participants from Jackson, Mississippi. The second exam (2000-2005) re-recruited 80% of participants. The GENOA data consists of biological samples (DNA, serum, urine) as well as demographic, anthropometric, environmental, clinical, biochemical, physiological, and genetic data. This study included 1157 African American participants.

*GOLDN*

Details of the TOPMed Project: Genetics of Lipid Lowering Drugs and Diet Network project can be found at: https://topmed.nhlbi.nih.gov/

*HyperGEN*

Details of the TOPMed Project: Hypertension Genetic Epidemiology Network and Genetic Epidemiology Network of Arteriopathy study can be found at: https://topmed.nhlbi.nih.gov/

*JHS*

The Jackson Heart Study (JHS, https://www.jacksonheartstudy.org/jhsinfo/) is a large, community-based, observational study whose participants were recruited from urban and rural areas of the three counties (Hinds, Madison and Rankin) that make up the Jackson, MS metropolitan statistical area (MSA). Participants were enrolled from each of 4 recruitment pools: random, 17%; volunteer, 30%; currently enrolled in the Atherosclerosis Risk in Communities (ARIC) Study, 31% and secondary family members, 22%. Recruitment was limited to non-institutionalized adult African Americans 35-84 years old, except in a nested family cohort where those 21 to 34 years of age were also eligible. The final cohort of 5,306 participants included 6.59% of all African American Jackson MSA residents aged 35-84 during the baseline exam (N-76,426, US Census 2000). Among these, approximately 3,700 gave consent that allows genetic research and deposition of data into dbGaP, with 3406 participants with post-QC TOPMed whole genome sequencing data. Major components of three clinic examinations (Exam 1 – 2000-2004; Exam 2 – 2005-2008; Exam 3 – 2009-2013) include medical history, physical examination, blood/urine analytes and interview questions on areas such as: physical activity; stress, coping and spirituality; racism and discrimination; socioeconomic position; and access to health care. Extensive clinical phenotyping includes anthropometrics, electrocardiography, carotid ultrasound, ankle-brachial blood pressure index, echocardiography, CT chest and abdomen for coronary and aortic calcification, liver fat, and subcutaneous and visceral fat measurement, and cardiac MRI. At 12-month intervals after the baseline clinic visit (Exam 1), participants have been contacted by telephone to: update information; confirm vital statistics; document interim medical events, hospitalizations, and functional status; and obtain additional sociocultural information. Questions about medical events, symptoms of cardiovascular disease and functional status are repeated annually. Ongoing cohort surveillance includes abstraction of medical records and death certificates for relevant International Classification of Diseases (ICD) codes and adjudication of nonfatal events and deaths. CMS data are currently being incorporated into the dataset.

*MESA*

The MESA study is a study of the characteristics of subclinical cardiovascular disease (disease detected non-invasively before it has produced clinical signs and symptoms) and the risk factors that predict progression to clinically overt cardiovascular disease or progression of the subclinical disease 7. MESA researchers study a diverse, population-based sample of 6,814 asymptomatic men and women aged 45-84. Thirty-eight percent of the recruited participants are white, 28 percent African-American, 22 percent Hispanic, and 12 percent Asian, predominantly of Chinese descent. Participants were recruited from six field centers across the United States: Wake Forest University, Columbia University, Johns Hopkins University, University of Minnesota, Northwestern University and the University of California - Los Angeles.

*VU_AF*

The Vanderbilt Atrial Fibrillation (AF) Registry was founded in 2001. Patients with AF and family members are prospectively enrolled. At enrollment a detailed past medical history is obtained along with an AF symptom severity assessment. Blood samples are obtained for DNA extraction. Patients are followed longitudinally along with serial collection of AF symptom severity assessments.

*WGHS*

The Women's Genome Health Study (WGHS) is a prospective cohort comprised of over 25,000 initially healthy female health professionals enrolled in the Women's Health Study, which began in 1992-1994. All participants in WGHS provided baseline blood samples and extensive survey data. Women who reported atrial fibrillation during the course of the study were asked to report diagnoses of AF at baseline, 48 months, and then annually thereafter. Participants enrolled in the continued observational follow-up who reported an incident AF event on at least one yearly questionnaire were sent an additional questionnaire to confirm the episode and to collect additional information. They were also asked for permission to review their medical records, particularly available ECGs, rhythm strips, 24-hour ECGs, and information on cardiac structure and function. For all deceased participants who reported AF during the trial and extended follow-up period, family members were contacted to obtain consent and additional relevant information. An end-point committee of physicians reviewed medical records for reported events according to predefined criteria. An incident AF event was confirmed if there was ECG evidence of AF or if a medical report clearly indicated a personal history of AF. The earliest date in the medical records when documentation was believed to have occurred was set as the date of onset of AF.

*WHI*

The Women’s Health Initiative (WHI) is a long-term, prospective, multi-center cohort study that investigates post-menopausal women’s health. WHI was funded by the National Institutes of Health and the National Heart, Lung, and Blood Institute to study strategies to prevent heart disease, breast cancer, colon cancer, and osteoporotic fractures in women 50-79 years of age. WHI involves 161,808 women recruited between 1993 and 1998 at 40 centers across the US. The study consists of two parts: the WHI Clinical Trial which was a randomized clinical trial of hormone therapy, dietary modification, and calcium/Vitamin D supplementation, and the WHI Observational Study, which focused on many of the inequities in women’s health research and provided practical information about the incidence, risk factors, and interventions related to heart disease, cancer, and osteoporotic fractures.

**Supplementary Methods**

Comparison of mCA calls from TOPMed WGS data and array-based calls in WHI

In total, there were 837 samples in the WHI with mCA calls from both TOPMed WGS as well as array-based data. To ensure comparability of the mCA callsets, we only considered samples where the DNA was isolated for TOPMed within 3 years of when the DNA was isolated for the array-based genotyping. We implemented the standard MoCha-based^1^ mCA calling detection pipeline using MoCha v1.14.

Array data from the WHI cohort, generated on hg19, were first lifted over to hg38, and then phased with shapeit4 (v4.2.2) against 1000 Genomes. Similar to the WGS, the MoChA caller was run with the extra option “--LRR-weight 0.0 --bdev-LRR-BAF 6.0” to disable the LRR+BAF mode. We also used the author-recommended flag “regression="--adjust-BAF-LRR -1 --regress-BAF-LRR -1" to avoid batch effects when processing array data. Given the unbalanced nature of gender in the WHI, we also provided this information directly using the option “--input-stats”. Subsequent calls were then excluded similarly to WGS: 1) those that span less than 500 informative sites, 2) those with estimated relative coverage higher than 2.9, and 3) those with BAF deviation larger than 0.16 and relative coverage higher than 2.5. Overlapping calls were ascertained using bedtools (v2.30.0). Disjoint calls (for example, if two calls on the same chromosome in one method overlap a single call in the second) were merged for tabulation purposes.

Of the 264 total MoCha-based array calls, 221 (83.7%) were found in the WGS data. On the other hand, WGS identified a total of 295 events, 221 (74.9%) of which were found in the array data. Overall concordance did not improve when higher clonal fractions (109 of 132 calls, 82.6%) were considered. We note that one of the primary drivers of the increased sensitivity in the WGS appears to be lower clonal fraction events (**Figures S7** & **S8**).

Burden Plots and Cholesky Residuals

**Figure S9** displays burden plots of Cholesky Residuals for each variant included in the aggregate rare-variant test for a number of different genes. The “working residuals'' are the difference of the “working outcome” vector and the linear predictor values from the final iteration of the penalized quasi-likelihood (PQL) approach used by the GMMAT^2^ method for solving the null logistic mixed model. The Cholesky residuals are a transformation of these working residuals computed using the estimated model covariance structure to remove the correlation. These Cholesky residuals are used to compute the variant scores that contribute to the association signals. For a burden test, positive residuals for case carriers are evidence for association between the rare allele at the variant and case status, while negative residuals for control carriers are evidence for association between the rare allele at the variant and control status. The magnitudes of the Cholesky residuals indicate the strength of evidence for association provided by that variant for that carrier, after adjustment for covariates and relatedness. The burden association signal for a gene aggregation unit depends on the summation of the Cholesky residuals for all of the carriers across all of the variants.
