## Supplementary material for "Mosaic chromosomal alterations in blood across ancestries via whole-genome sequencing": Figures S1-S8

**Supplementary Figure 1: Distribution of mCA mutant cell fraction estimates**

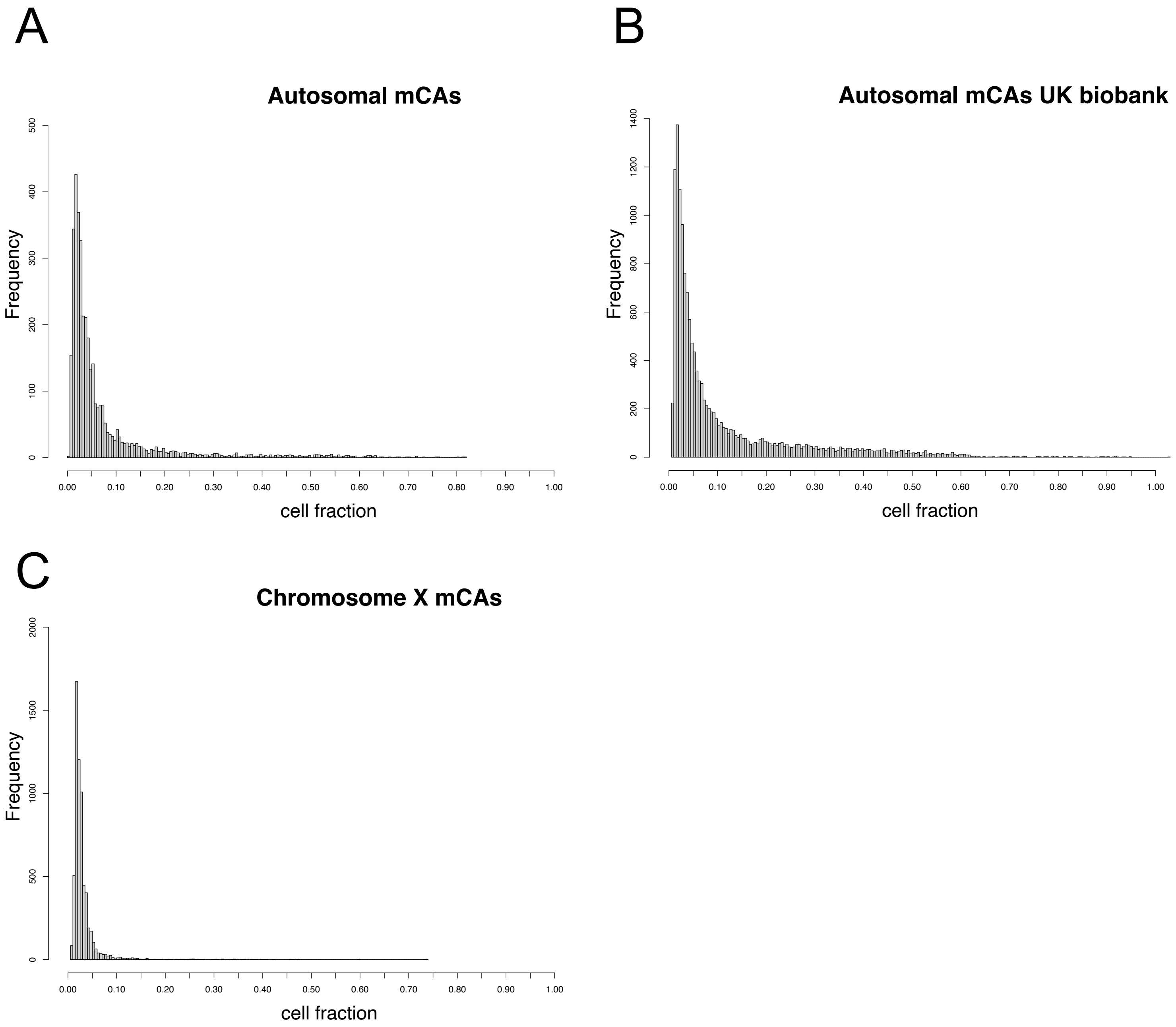


Histogram of cell fraction estimates for A) autosomal mCAs in males and females B) autosomal mCAs in UKBB C) chromosome X mCAs in females

**Supplementary Figure 2: Genomic distribution of mCA across mutant cell fraction categories**

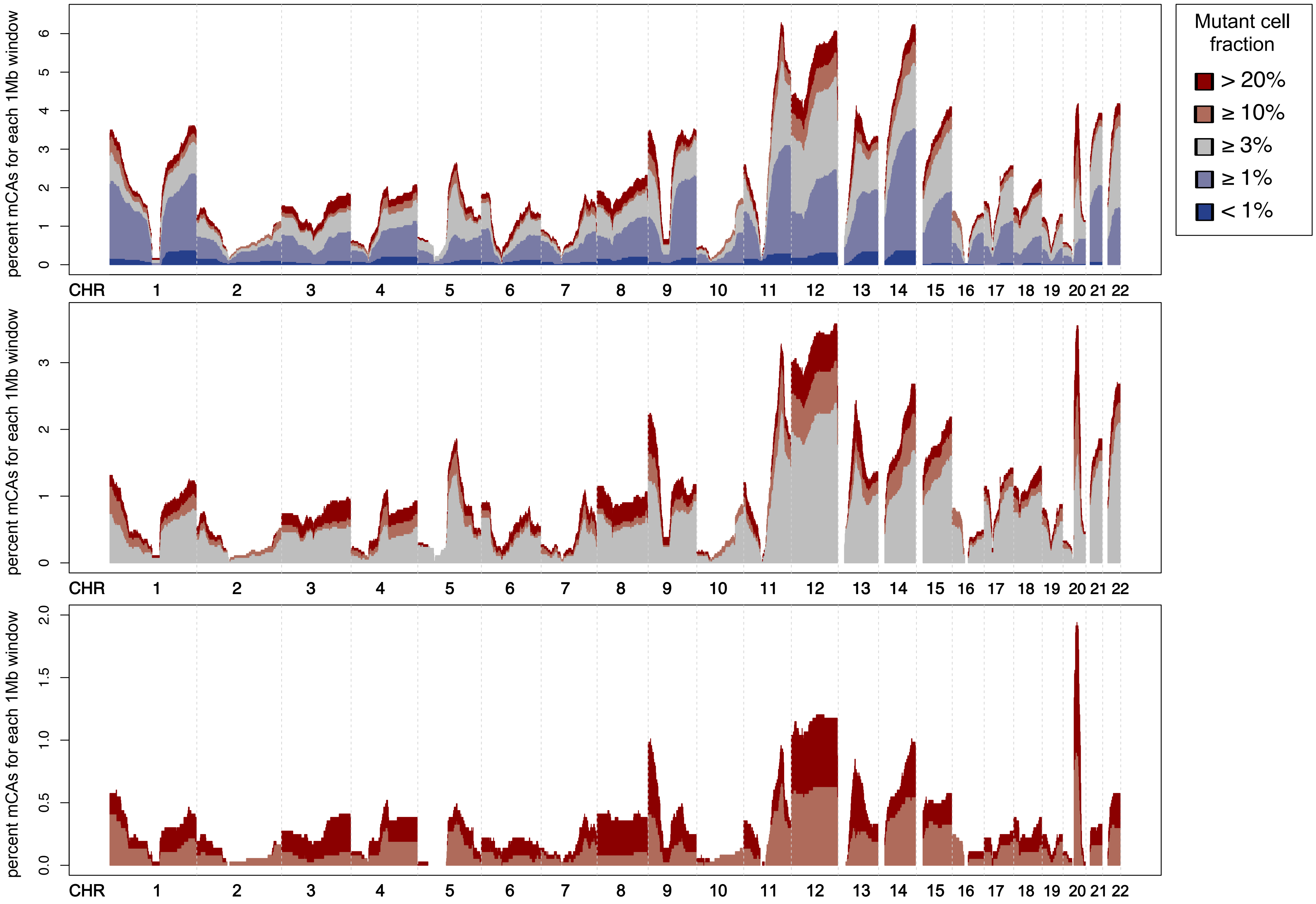


Histogram with the percent of mCAs (number of mCA calls spanning genomic location / all mCA calls). Bins span 1Mb.

**Supplementary Figure 3: Genomic distribution of mCA across AA and EA ancestries for different mutant cell fraction categories**

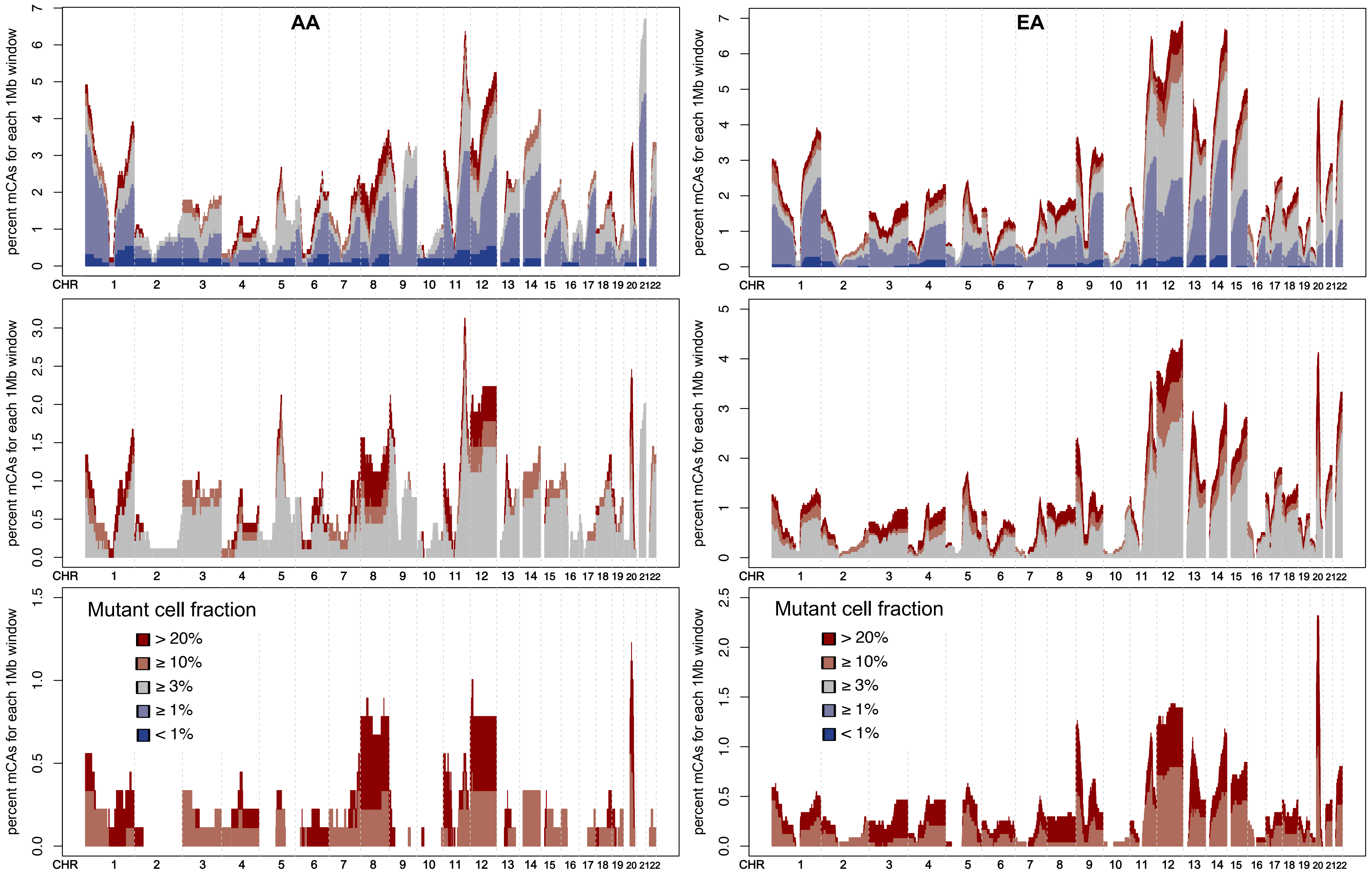


Histograms with percent of mCA calls per genomic region. Bins span 1Mb.The 3 histograms on the left are for mCAs detected in individuals of African American Ancestry (AA) and the three histograms on the right are for mCAs detected in individuals of European Ancestry (EA).

**Supplementary Figure 4: Rate mCAs by age in females**

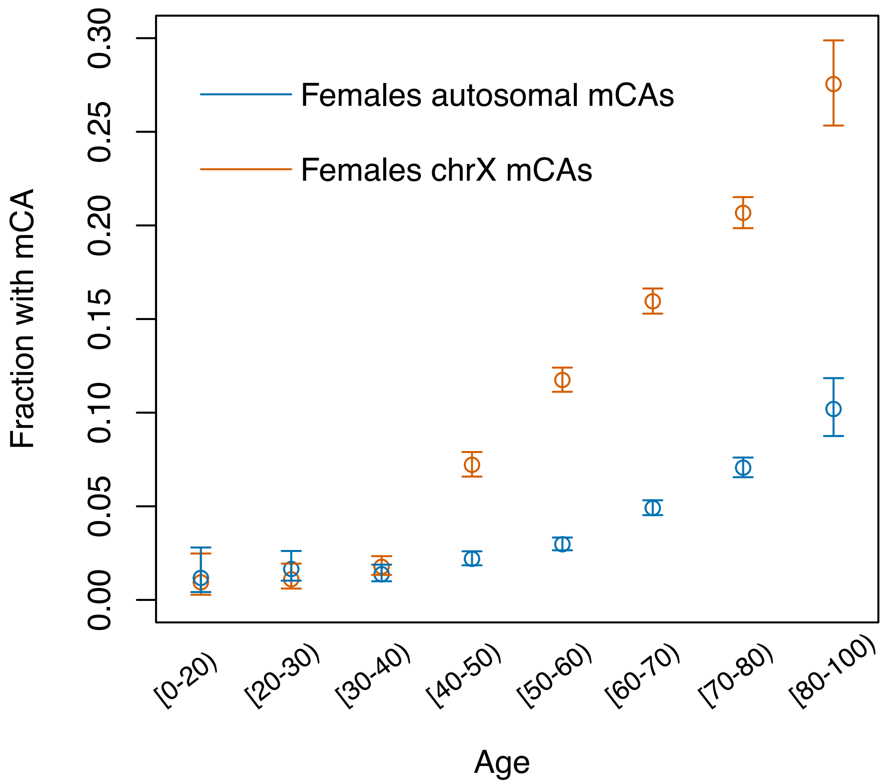


**Supplementary Figure 5: Rates chrX mCAs by genetic ancestry**

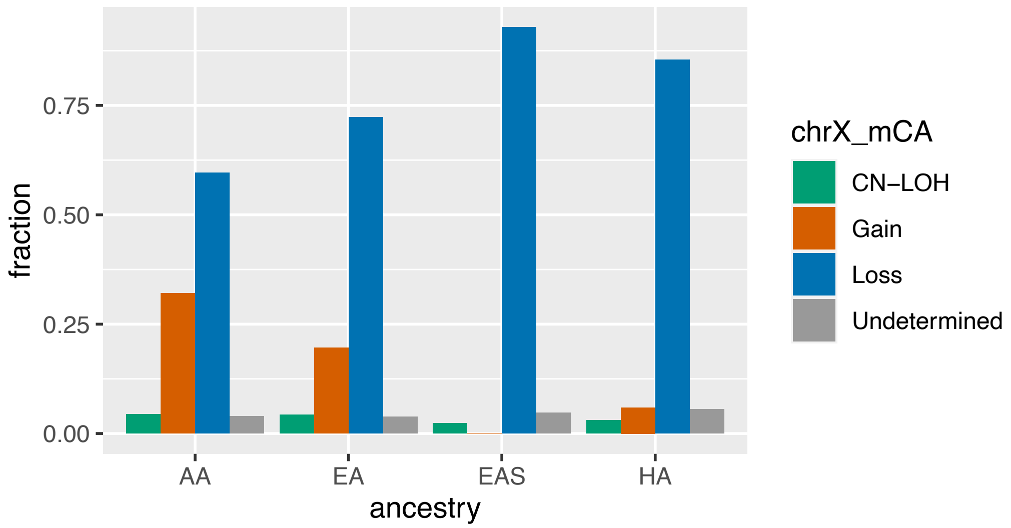

**Supplementary Figure 6:**

A

**lymphoid_cancer**

| mCA_auto carrier | Control (Yes/No) | Case (Yes/No) | OR (mean [95% CI]) |
|---|---|---|---|
| All | 441/7360 | 32/185 | 2.96 [1.9,4.44] |
| Not cytopenia | 382/6194 | 20/152 | 2.41 [1.43,3.85] |
| Not cytosis | 416/7145 | 21/180 | 1.98 [1.14,3.21] |
| Not cytopenia either non-cytosis | 362/6051 | 14/149 | 1.74 [0.93,3] |

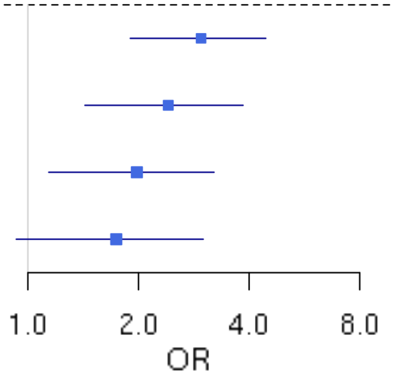


**B**

**myeloid_cancer**

| mCA_auto carrier | Control (Yes/No) | Case (Yes/No) | OR (mean [95% CI]) |
|---|---|---|---|
| All | 441/7360 | 10/42 | 5.42 [2.39,11.17] |
| Not cytopenia | 382/6194 | 5/33 | 3.68 [1.22,9.11] |
| Not cytosis | 416/7145 | 7/39 | 4.04 [1.49,9.26] |
| Not cytopenia either non-cytosis | 316/5365 | 4/26 | 4.06 [1.15,11.16] |

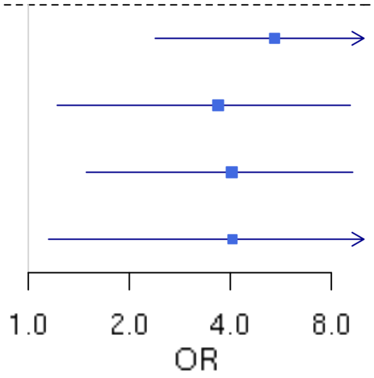

**Supplementary Figure 7: Histogram of mCA calls across clonal fractions**

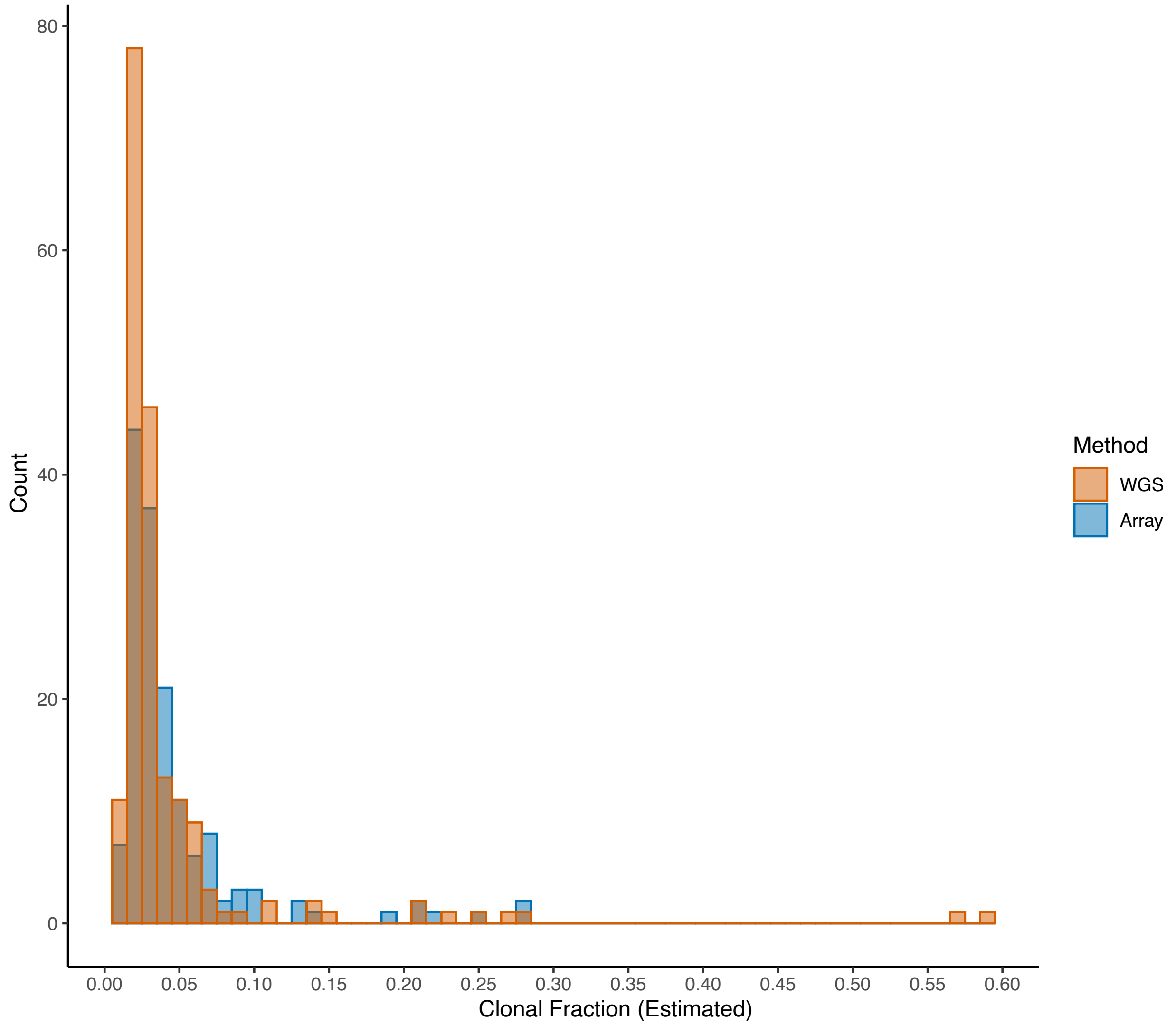


Lower clonal fraction mCAs are detected with higher sensitivity with WGS data compared to array data. Calls made from either the WGS- and array-data are plotted.

**Supplementary Figure 8: Histogram of mCA calls across clonal fractions**

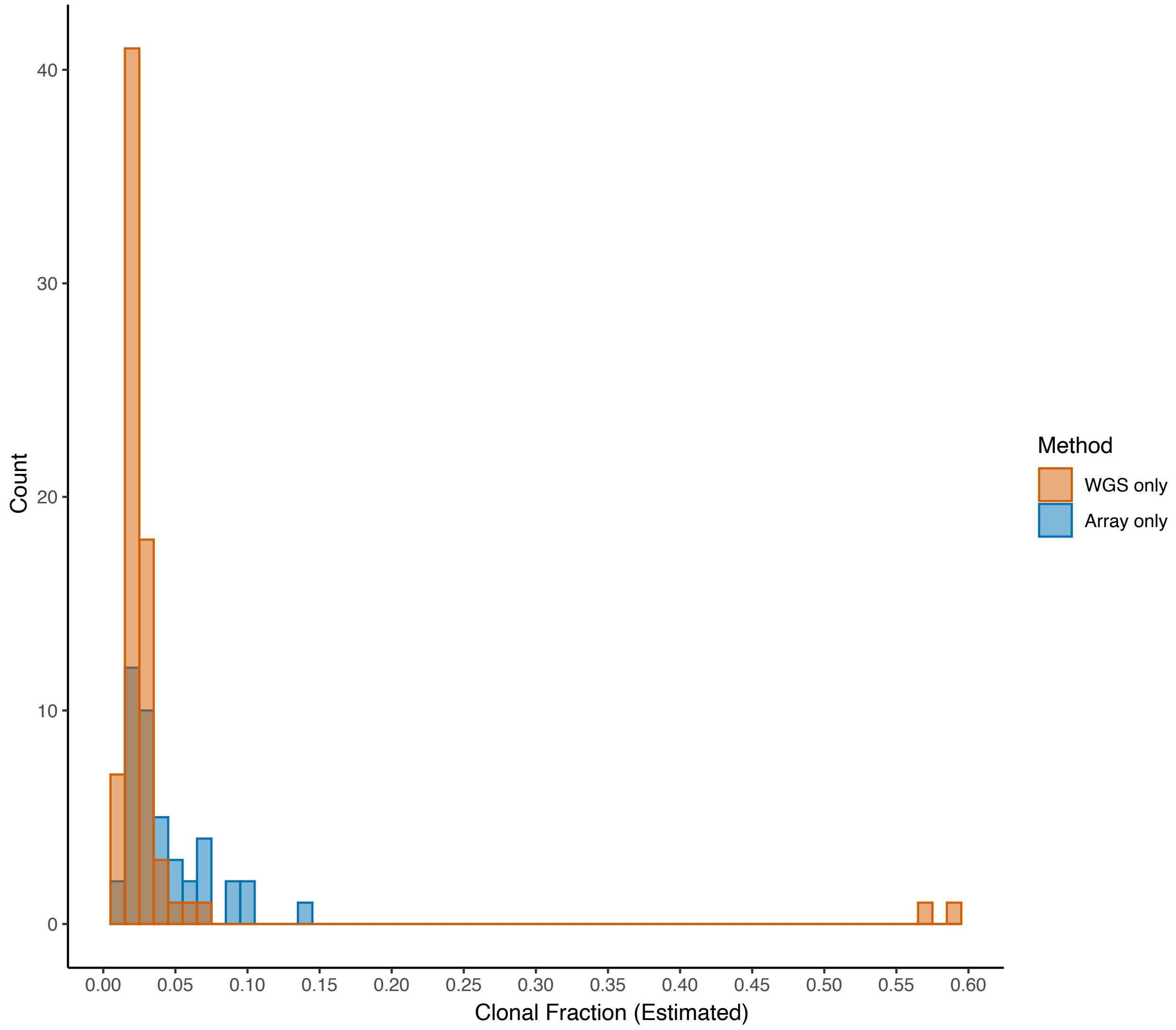


Lower clonal fraction mCAs are detected with higher sensitivity with WGS data compared to array data. Calls made from both the WGS- and array-data are plotted.
